# Deconvolution of Sparse-count RNA Sequencing Data for Tumor Cells Using Embedded Negative Binomial Distributions

**DOI:** 10.1101/2025.11.21.689822

**Authors:** Matthew D Montierth, Hao Yan, Liyang Xie, Kinga Nemeth, Xiaoxi Pan, Ruonan Li, Caner Ercan, Peng Yang, Ansam Sinjab, Tieling Zhou, Fuduan Peng, Manisha Singh, Linghua Wang, Scott Kopetz, Humam Kadara, Yinyin Yuan, George A. Calin, Wenyi Wang

## Abstract

Estimating tumor-specific transcript proportions from mixed bulk samples has potential to inform novel biology. However, estimation accuracy using existing methods in sparse-count data such as microRNA-seq and spatial transcriptomics has yet to be established. We generated a mixed small RNA benchmark dataset to demonstrate analytical challenges. To resolve them, we developed DeMixNB, a semi-reference-based deconvolution model assuming a sum of negative binomial distributions. Applications to miRNA-seq from 856 patients with breast cancer and 3,755 spatial spots from lung cancer generated either clinical or mechanistic insights into tumor cell plasticity. This supports the important utility of DeMixNB to investigate cancer RNomes.

## Introduction

Cancer accounts for nearly one in six deaths globally and is a leading cause of mortality across both developed and developing nations^1^. Despite decades of research and improvements in therapies, treatment outcomes vary, reflecting not only differences in access and care but also profound biological variation between tumors^2^. Cancer’s complexity partly stems from the presence of intratumor heterogeneity and the dynamic nature of tumor cell plasticity^3,4^. Variability within a tumor arises from the complex cellular composition in the local tumor microenvironment (TME), impacting progression, spread and therapeutic resistance^5,6^. Adding to this cellular complexity, the plasticity of malignant cells, which is the ability of cancer cells to transit between distinct stages of differentiation and a state of ‘stemming’, fosters the tumor diversity^7^. Such plasticity-driven variations significantly influence the total amount of transcripts originating from tumor cells^8^. Previous studies^9–11^ showed that tumor cells exhibit much greater transcriptional activity and variation compared to surrounding non-tumor cells. Therefore, accurately quantifying tumor-cell specific transcript proportion of tissues offers direct insights into tumor heterogeneity and is becoming a unique and desirable feature to obtain for cancer studies.

One common computational strategy to perform the aforementioned task is deconvolution, a mathematical technique that can separate mixed signals from tissues, enabling the estimation of cell-type proportions, for either cell counts or cell transcripts, and cell specific transcript proportion to reveal tumor heterogeneity^12^. The most widely used deconvolution framework is reference-based deconvolution techniques require finding marker genes with stably expressed profiles for all cell types of interest, hence not suitable to capture the yet-to-measure transcriptional plasticity in tumor cells. Reference-free methods^13–18^, on the contrary, make minimal assumptions on the cell-type specific gene expression profiles, but consequently cannot achieve as good estimation performances. Semi-reference-based deconvolution^19–28^ provides a middle ground, requiring reference data from cell types that are relatively stable expressed while offering flexibility on the tumor-cell gene expression. This setup makes semi-reference-based best suited for estimating tumor-specific transcript proportions.

Existing semi-reference-methods like DeMix^24,25^/DeMixT and ISOpure^29^ use expression profiles of adjacent normal tissues as the partial reference. Other methods like PREDE^21^ and BayICE^28^ rely on Non-negative Matrix Factorization (NMF) and Markov chain Monte Carlo procedure through Gibbs sampling pre-selected cell-type marker genes and number of optimal cell types to perform deconvolution. By integrating partial reference data, semi-reference methods can address the issues of instability of deconvolution results, incomplete cellular knowledge and technical variability. While these methods provide a versatile and reliable solution for dissecting the TME’s mixed cellular compositions^12^, they were primarily developed for high read count data like bulk RNA sequencing (RNA-seq) data. For example, the log normal distribution assumption in DeMixT is strongly violated when genes present low read counts. This then limits the application of DeMixT to the deconvolution of several important RNomes, such as microRNA sequencing^30^ (miRNA-seq) or spot-level spatial transcriptomic (ST) data^31^.

MiRNAs and other non-coding RNAs are extensively linked to biology and disease pathogenesis^32^, although no single-cell sequencing technology has been established for miRNA. Spatial transcriptomics from popular technologies such as 10X Genomics Visium faces two major analytical challenges: sparsity from low capture efficiency and mixed signals from each spot capturing 5-50 cells of multiple cell types. Reference-based deconvolution methods have been developed recently for both bulk miRNA-seq (DeconmiR^33^), and ST (RCTD^34^, CARD^35^ data. Still, no semi-reference-based methods have been developed to study tumor-cell transcript proportions in these data types specifically.

To fill this gap, we have developed DeMixNB that models the tumor and non-tumor cell compartments as individual negative binomial (NB) distributions for low read counts and takes the reference profiles from the non-tumor cell compartment only as input (**Figure 1**). Using our specially designed cell-line mixing miRNA experiment as a benchmark dataset for low read count data, we found DeMixNB to significantly outperform existing methods in estimating tumor-cell transcript proportions. We further provide two real data applications to demonstrate that for both miRNA-seq and ST Visium data, tumor-cell transcript proportions are distinct from tumor-cell count proportions, i.e., purity, and have the potential to inform prognosis-associated transcriptional activities in tumor cells as well as their interaction with TME. DeMixNB is freely available within the R package DeMixT 2.0.

**Figure 1.**
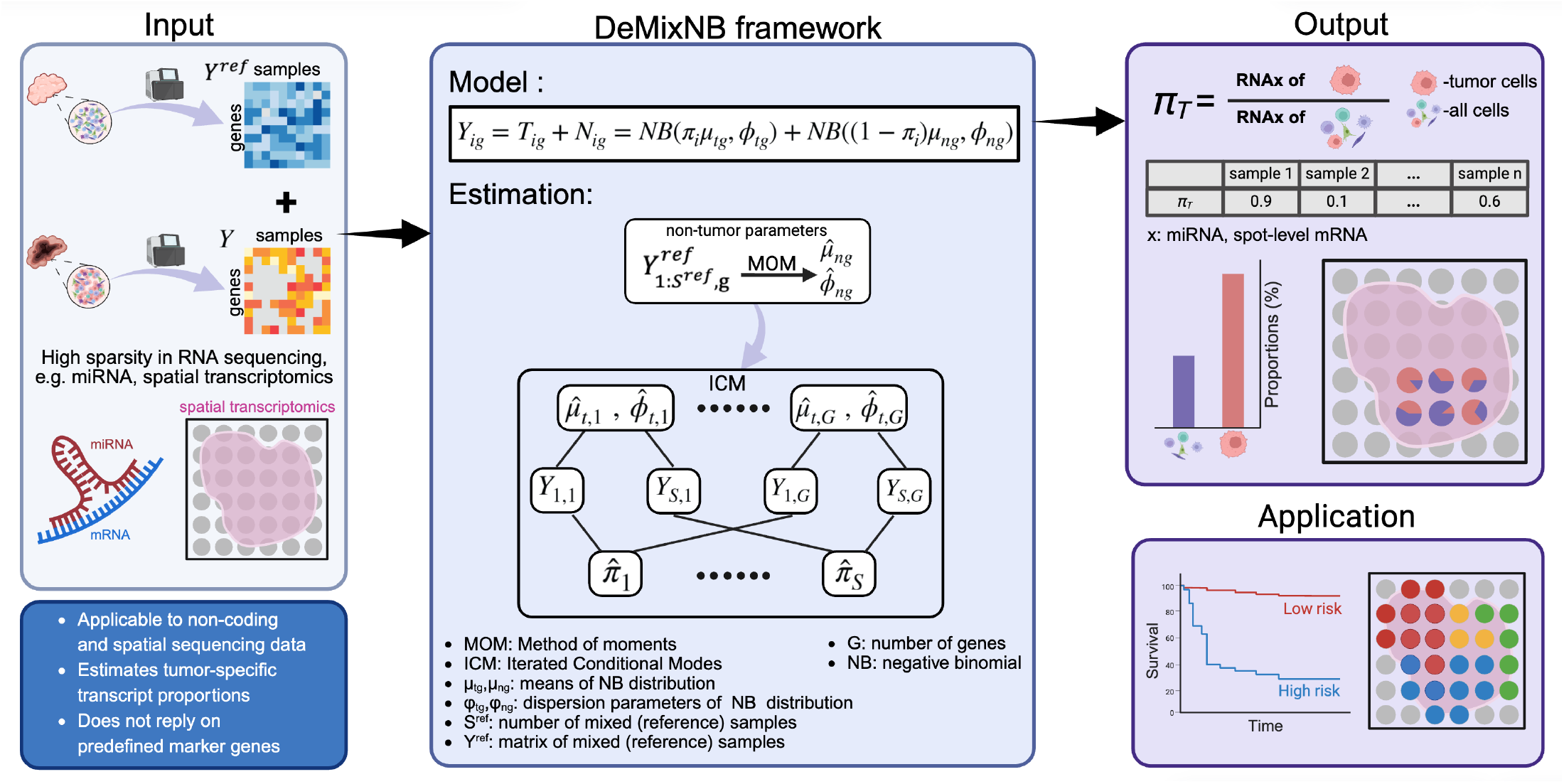
Overview of DeMixNB. DeMixNB takes gene expression profiles from mixed samples and corresponding reference samples as input (sample-by-gene for bulk sequencing data or spot-by-gene for spatial transcriptomic data). DeMixNB estimates tumor-specific transcript proportions without the need for external reference or cell type marker gene list. Observed gene expression *Y*_*ig*_ is modeled as the sum of reference component *N*_*ig*_ and tumor component *T*_*ig*_, with each following a negative binomial distribution respectively. Tumor-transcript proportion *π*_*i*_ is estimated simultaneously with the mean *µ*_*ig*_ and overdispersion parameter ∅_*ig*_. For each mixed sample, DeMixNB estimates *π*_*T*_, which is the tumor transcript proportion of the RNA species being studied. In spatial transcriptomics, this is estimated for each spot, which can be used to refine profiling of spatial patterns in tumor tissues. For sparse non-coding RNAs like miRNAs, the *π*_*T*_-miR proportion can be used as a biological feature in downstream analyses. Similar application is also feasible for the spot-level *π*_*T*_.

## Results

### DeMixNB overview

We propose DeMixNB, a negative-binomial-based semi-reference-free deconvolution method that is designed for low-abundance (sparse) transcriptomic data. Previously, methods for spatial transcriptomics have attempted to address the sparsity problem through the use of negative binomial distributions, which is inherently better suited for sparse count data^31^. However, these methods require extensive single-cell reference data or prior marker gene knowledge, which results in complex model structures with hyperparameters to adjust for the technological differences and batch effects, making them unsuitable for use cases without single-cell reference. Another drawback with these methods is that they estimate cell-type proportions instead of transcript proportions, making them less useful in understanding tumor-specific transcript activity. DeMixNB addresses these challenges by combining the semi-reference approach of DeMixT^24^, which requires only reference samples of gene expression levels in non-tumor cells, with negative binomial modeling optimized for sparse count data, such as those from miRNA-seq and ST (**Supplementary Figures S1-2**).

The observed expression for each gene *g* and sample *i, Y*_*ig*_ is assumed to be the sum of expressions in tumor cells, *T*_*ig*_, and expressions in non-tumor cells, *N*_*ig*_. Both *T*_*ig*_and *N*_*ig*_ are assumed to follow negative binomial distributions specified by the mean and overdispersion parameters *µ*_*tg*_, *ϕ*_*tg*_, and *µ*_*ng*_, *ϕ*_*ng*_ respectively. The proportion parameter, *π*_*i*_, ensures the mean of mixed expressions *E*(*Y*) = *π µ* + (1 − *π*)*µ*, whose estimate approximates the transcript proportion 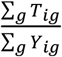 (see further **in Methods**). Given an input expression matrix representing mixtures of tumor and non-tumor cells and a reference expression matrix representing only non-tumor cells, DeMixNB first estimates parameters for the non-tumor component 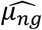 and 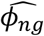, then performs the iterative conditional modes (ICM)^36^ to efficiently optimize the likelihood of the target mixed samples: iteratively estimating the sample-wise parameters - the mixing proportions of tumor samples, and then the gene-wise parameters - the distribution parameters of individual gene expressions. The ICM approach is used to address an intrinsic weak tractability of all model parameters in the model’s full likelihood. Our key assumption is that the proportions only affect the mean of the distributions of expressions, while the dispersion parameters remain independent of proportions, in order to achieve computational tractability (**Methods**).

### Simulation study

To verify the convergence of the ICM approach in DeMixNB and the estimation accuracy, we introduced multiple setups of simulated data with known ground truth transcript compositions. We considered three simulation scenarios which differ by level of sparsity and gene expression distributions: A. non-sparse count data, B. sparse count data with a single tumor-specific distribution per gene, C. sparse count data with two tumor-specific distributions across genes representing tumor cell subtypes across samples (**Figure 2a, Methods**). We benchmarked the DeMixNB performance against DeMixT^24^, PREDE^21^ and ISOpure^23^, three established semi-reference-based deconvolution methods, by comparing the estimated proportions produced by each model with the true tumor transcript proportions 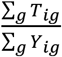. Our evaluation metrics include the absolute bias, concordance correlation coefficient (CCC), the root mean square error (RMSE), and the consistency metric (sumEst) that sums up the proportion estimates using the nontumor and tumor reference respectively, hence a perfect score is 1. The resulting metrics and p-values for paired-t-tests are summarized in **Table 1** and visualized in **Figure 2**.

**Table 1.**
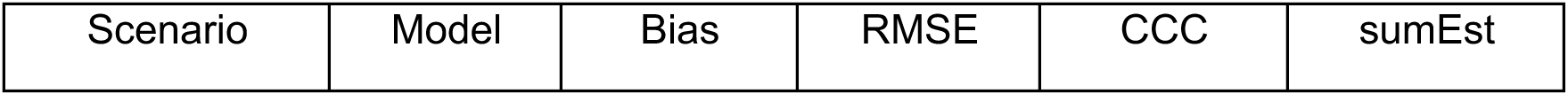
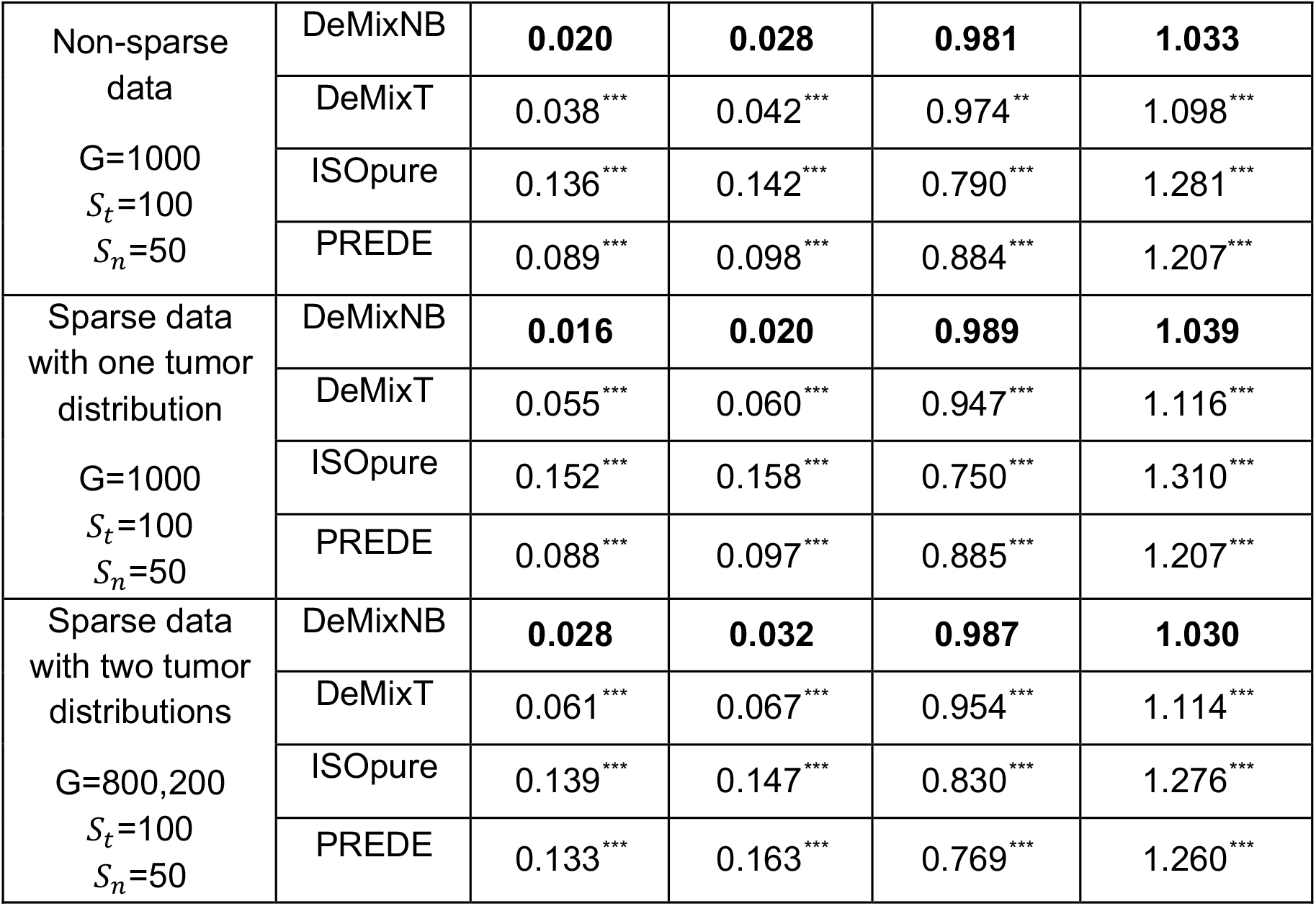
Simulation study. The metric sumEst represents the sum of the estimated tumor proportions using non-tumor reference and the estimated non-tumor proportions using tumor reference. Paired-t-tests have been conducted between DeMixNB and the other three methods and the Benjamini-Hochberg-adjusted-p-values for each metric are compared with different alpha levels and reported in the table (**: adjusted-p < 0.001, **: adjusted-p < 0.01, *: adjusted-p < 0.05). The best performing model for each metric is bolded.

| Scenario | Model | Bias | RMSE | CCC | sumEst |
| --- | --- | --- | --- | --- | --- |
| Non-sparse data<br>G=1000<br>$S_t=100$<br>$S_n=50$ | DeMixNB | **0.020** | **0.028** | **0.981** | **1.033** |
|  | DeMixT | 0.038*** | 0.042*** | 0.974** | 1.098*** |
|  | ISOpure | 0.136*** | 0.142*** | 0.790*** | 1.281*** |
|  | PREDE | 0.089*** | 0.098*** | 0.884*** | 1.207*** |
| Sparse data with one tumor distribution<br>G=1000<br>$S_t=100$<br>$S_n=50$ | DeMixNB | **0.016** | **0.020** | **0.989** | **1.039** |
|  | DeMixT | 0.055*** | 0.060*** | 0.947*** | 1.116*** |
|  | ISOpure | 0.152*** | 0.158*** | 0.750*** | 1.310*** |
|  | PREDE | 0.088*** | 0.097*** | 0.885*** | 1.207*** |
| Sparse data with two tumor distributions<br>G=800,200<br>$S_t=100$<br>$S_n=50$ | DeMixNB | **0.028** | **0.032** | **0.987** | **1.030** |
|  | DeMixT | 0.061*** | 0.067*** | 0.954*** | 1.114*** |
|  | ISOpure | 0.139*** | 0.147*** | 0.830*** | 1.276*** |
|  | PREDE | 0.133*** | 0.163*** | 0.769*** | 1.260*** |

**Figure 2.**
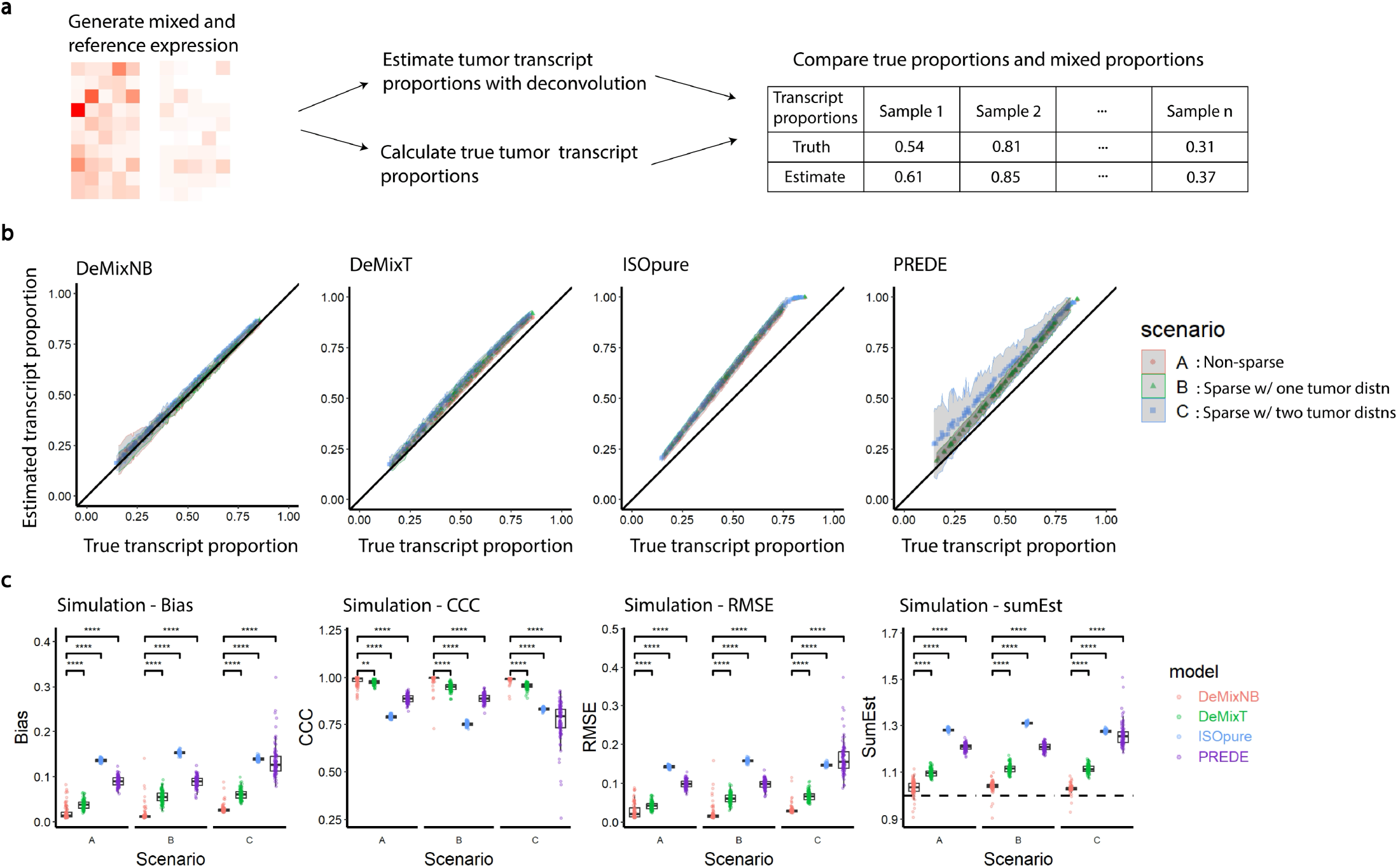
Simulation study and results. (a) An overview of the simulation process. (b) Scatter plots with confidence ribbons (mean ± standard error) for the estimated transcript proportions output by each deconvolution method (y-axis) and the true transcript proportion calculated from the generated data (x-axis). Simulation settings are marked with different colors. See **Methods** for more details. (c) Boxplots comparing the average Bias, CCC, RMSE, and sum of estimated proportions (sumEst) over all 100 mixed samples in each of the 100 simulated datasets across all simulation scenarios for DeMixNB, DeMixT, ISOpure, and PREDE. For Bias and RMSE, the smaller the better. For CCC, the closer to 1, the better. The sumEst metric calculates the sum of the estimated tumor proportion using non-tumor reference and the estimated non-tumor proportions using tumor reference. The closer it is to 1, the more consistent we consider the model to be. All result comparisons with DeMixNB are significant as shown by the non-overlapping quartile ranges and further supported by pairwise t tests (significance level denoted by: *, p < 0.05; **, p < 0.01; ***, p < 0.001; ****, p < 0.0001.

Across all three scenarios, DeMixNB displayed consistently better performance in specifically estimating transcript proportions with significantly higher CCC, lower Bias and RMSE, and more consistency as measured by sumEst (**Figure 2b-c**). Both DeMixNB and DeMixT did well with nonsparse data while in this case DeMixT showed a computational speed advantage by assuming a continuous log-normal distribution. When applied to the simulated sparse data, DeMixT, ISOpure, and PREDE all tend to overestimate the tumor transcript proportions (**Figure 2b**). As the data becomes sparser in scenario B compared to scenario A, both DeMixT and ISOpure displayed a slight drop in performance while DeMixNB maintained similarly good performance in terms of RMSE and CCC, and the results for PREDE remained the same. In scenario C with two different tumor distributions, the most notable drop in performance was observed for PREDE as its estimations became unstable across repeated runs. We also observed a slight drop in performance for DeMixNB and ISOpure while DeMixT results were similar to scenario B. In all settings, the CCC value for DeMixNB is above 0.98, representing a high accuracy in estimating transcript proportions, when the underlying gene expression distributions are negative binomial. Overall, the simulated data generated by the negative binomial distribution posed challenges to all methods except DeMixNB. Such results are expected since the normality assumption for DeMixT and PREDE is explicitly violated and the hyperparameter sampling used to generate these samples violates the assumption for ISOpure that the tumor samples belong to the same cancer subtype, such that the expressions are closely clustered cannot be satisfied.

### DeMixNB accurately deconvolves miRNA-seq in a specially-designed mixed cell line benchmarking study

To evaluate DeMixNB’s performance with sparse expression count matrices typical of bulk miRNA sequencing (miRNA-seq, **Supplementary Figure S1**), we developed a benchmarking dataset with precisely known cellular compositions (**Figure 3a**), a strategy that has been proven to be useful in benchmarking semi-reference based deconvolution methods^11^. The dataset comprised thirty-nine samples created by systematically mixing HS-5 bone marrow stromal fibroblast cells with either wild-type HCT116 colorectal cancer cells or CRISPR-Cas9-engineered *Dicer1* knockout HCT116 colorectal cancer cells (**Methods**). *Dicer1* encodes an enzyme required to process most precursor microRNAs into their mature forms^37^. Knocking out *Dicer1* causes global depression of mature miRNA levels^38^, which we confirmed via direct measurement of miRNA concentration per million cells (**Figure 3a**). To simulate diverse tumor microenvironment scenarios, we established seven distinct miRNA mixing ratios (0:100, 30:70, 50:50, 60:40, 70:30, 80:15, and 100:0 tumor:fibroblast), with three independent biological replicates generated for each ratio. Importantly, these mixed samples were generated after measuring the total miRNA density per million cells (**Figure 3a**), with extracted miRNA mixed to achieve the specified tumor-stromal total miRNA ratio, rather than cell count ratio. Furthermore, the tumor component across samples presents a large variation in total miRNA, representing either *Dicer1* wildtype or knockout cells. This design, featuring known ground truth proportions of both tumor-cell miRNA transcript and tumor-cell count and tumor cell plasticity, allowed us to thoroughly benchmark DeMixNB’s accuracy across a range of scenarios typically encountered in patient-derived samples.

**Figure 3.**
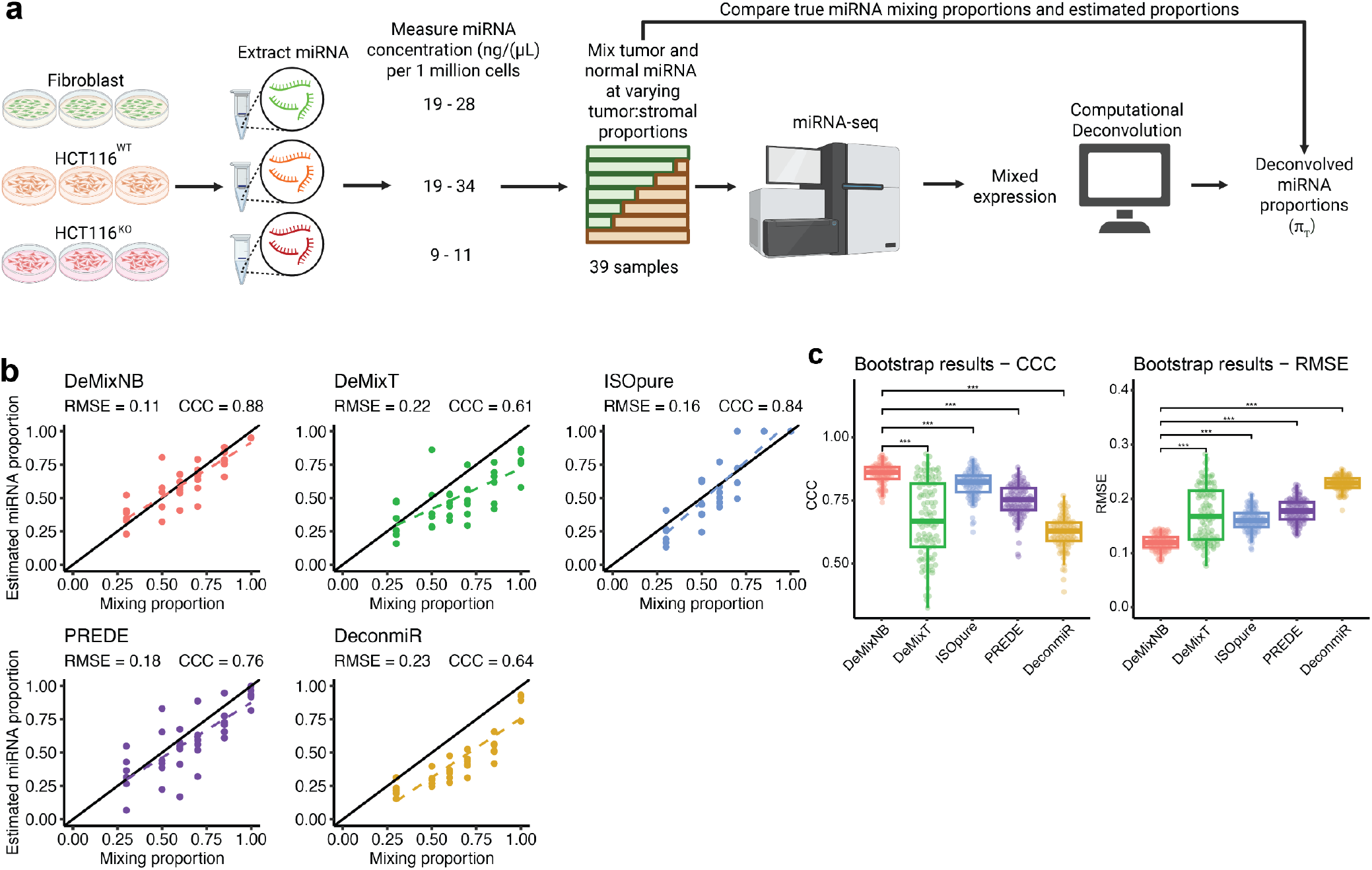
Benchmarking study design and results. (a) Graphical depiction of how the benchmarking dataset was generated **(Methods)**. (b) Scatter plots showing the relationship between tumor-specific total miRNA transcript proportion (x-axis) and the corresponding estimated proportion (y-axis) for five methods: DeMixNB, DeMixT, ISOpure, PREDE, and DeconmiR. Each plot includes a diagonal reference line (slope = 1) and a dotted line indicating linear regression fit. RMSE and CCC values are displayed on each plot. (c) Boxplots showing bootstrapped distributions of CCC and RMSE for DeMixNB, DeMixT, ISOpure, PREDE, and DeconmiR. Pairwise Wilcoxon rank-sum test p-values (BH adjusted) are indicated. For all plots, significance levels are denoted by: *, p < 0.05; **, p < 0.01; ***, p < 0.001

DeMixNB demonstrated high accuracy in predicting the true transcript proportions, achieving a root mean square error (RMSE) of 0.11 and a concordance correlation coefficient (CCC) of 0.88, with good accuracy observed across all mixing ratios (**Figure 3b, Methods**). When compared to existing semi-reference-based deconvolution methods on the same dataset, DeMixNB outperformed DeMixT (RMSE = 0.22, CCC = 0.61), ISOpure (RMSE = 0.16, CCC =0.84), and PREDE (RMSE = 0.18, CCC = 0.76), with consistently lower RMSE and higher CCC in bootstrapped resampling (adjusted pairwise Wilcoxon p-values < 0.001, **Figures 3b-c**). Additionally, all semi-reference-based methods outperformed DeconmiR, a reference-based miRNA deconvolution method (RMSE = 0.23, CCC = 0.64) (**Figure 3b, Supplementary Figures S3-4, Methods**), supporting the importance of making minimal assumptions on gene expression for the tumor-cell component. These results suggest that DeMixNB can reliably estimate cell-type specific transcript proportions from bulk miRNA-seq data with greater accuracy than current methods, even in complex mixed samples with highly variable tumor specific transcripts, i.e., between *Dicer1* wildtype and knockout cells, which are deconvolved together in one batch.

### Application to bulk miRNA-seq data from breast cancer

To demonstrate the utility of tumor-specific transcript proportions of miRNA in cancer, we applied DeMixNB to a miRNA-seq dataset from 856 patients with breast cancer (BCa) in The Cancer Genome Atlas projects (TCGA)^39^, where the non-tumor miRNA expression profiles can be derived from adjacent normal tissues of 98 patients (**Methods**). Prior work has established that miRNAs play key roles in breast cancer pathogenesis, where expression of oncogenic miRNAs (oncomiRs) and tumor-suppressive miRNAs contribute to disease progression and therapeutic resistance^32,40–44^. However, these studies have focused on individual miRNAs or on signatures built from groups of miRNAs, and as such the consequence of global tumor-cell miRNA abundance remains poorly defined. By quantifying tumor-cell miRNA proportion (*π*_*T*_-miR), DeMixNB provides a robust framework to examine this understudied domain of tumor microRNA biology.

The deconvolved tumor-cell total miRNAs proportion *π*_*T*_-miR varied significantly across samples and PAM50 molecular subtypes, which is a classification system that stratifies breast cancers into biologically distinct groups (Luminal A, Luminal B, HER2-enriched, and Basal-like; **Figure 4a**). Luminal A (LumA) breast cancers exhibited the lowest median *π*_*T*_-miR values and showed greater heterogeneity compared to other PAM50 subtypes, indicating that the tumor total miRNA proportion may be biologically informative in this subtype (**Figure 4a**). Recursive partitioning of LumA-BCa samples into high- and low *π*_*T*_-miR groups showed that patients with high *π*_*T*_-miR had significantly worse progression-free survival (PFS, log-rank p = 0.013, **Figure 4b, Methods**). This relationship persisted in a multivariate Cox regression after adjusting for clinical stage and patient age (**Figure 4c**), highlighting the independent prognostic value of *π*_*T*_-miR. Based on our simulation study, we chose to compare *π*_*T*_-miR with well-benchmarked DNA copy number-based tumor cell count proportion, i.e., tumor purity estimates, instead of DeconmiR. While *π*_*T*_-miRs showed a moderate positive correlation with tumor purity (Pearson correlation = 0.53), they were consistently higher than the corresponding purity values, suggesting an overall increased miRNA transcriptional activity within tumor cells as compared to nontumor cells (**Figure 4d**). Finally, the recursive partitioning of tumor purity within LumA-BCa samples fails to significantly distinguish PFS (**Figure 4e, Methods**), underscoring the biological and clinical information captured uniquely by *π*_*T*_-miRs, as estimated by DeMixNB.

**Figure 4.**
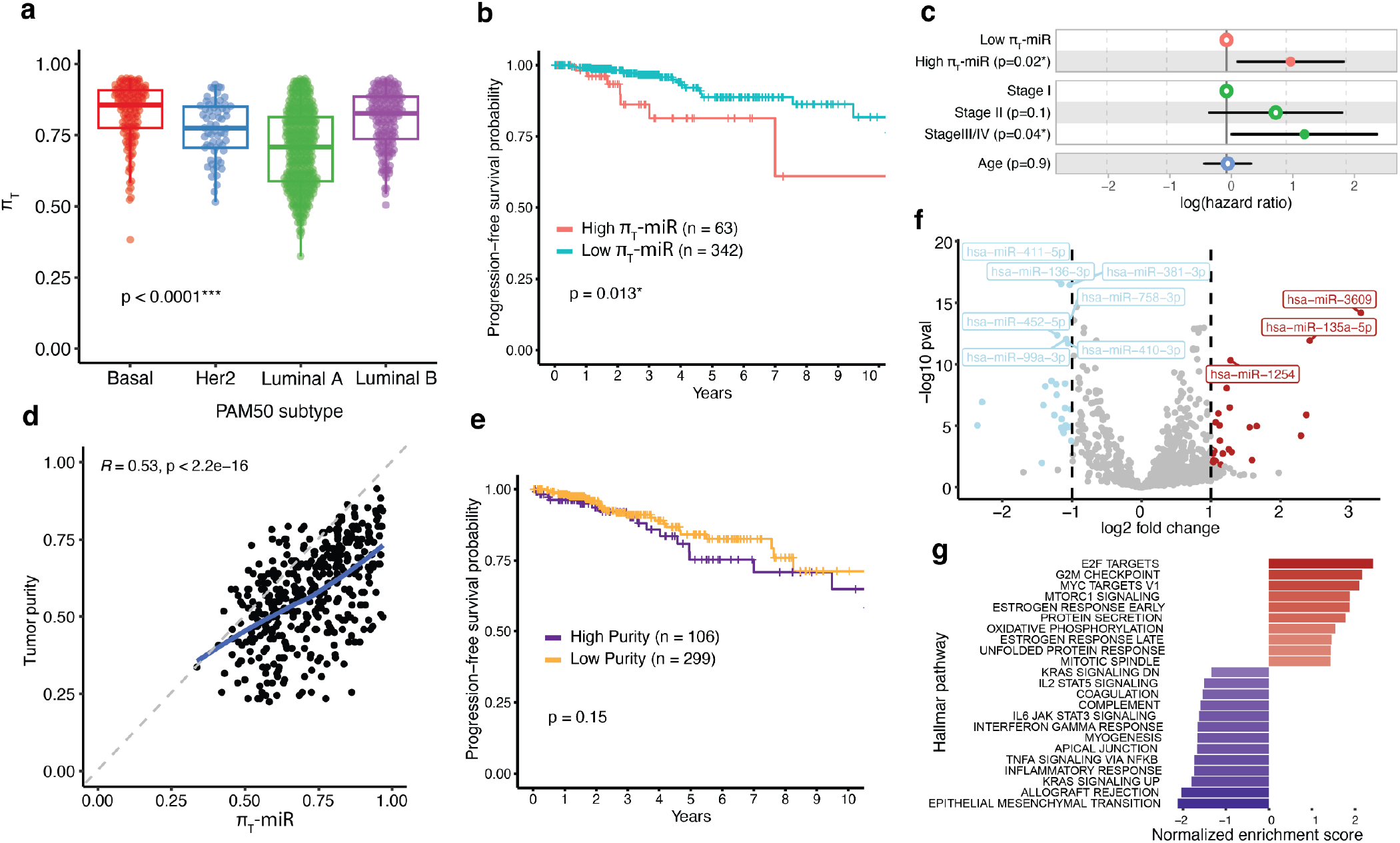
Deconvolution of bulk miRNA-seq for TCGA breast cancer. (a) Boxplots showing *π*_*T*_ estimates for TCGA breast cancer patients, separated by PAM50 subtype, with the Kruskal-Wallis p-value shown. (b) Kaplan-Meier curves showing progression-free survival within the Luminal A breast cancer (LumA-BCa) samples, stratified by *π*_*T*_ values (high vs. low, cutoff = 0.85), with the log-rank test p-value shown. (c) Forest plot showing hazard ratios for a Cox proportional hazard (PH) model fit on the LumA-BCa samples, with *π*_*T*_, stage, and age as covariates. Points represent the estimated hazard ratios, with 95% confidence intervals indicated by the lines. (d) Scatter plot showing relationship between *π*_*T*_ and tumor purity. The dashed gray line indicates slope = 1, with the blue line showing the lowess fit of the data. Pearson correlation coefficient is shown. (e) Kaplan-Meier curves showing progression-free survival within the LumA-BCa samples, stratified by purity values (high vs. low, cutoff = 0.66), with the log-rank test p-value shown. (f) Volcano plot of differentially expressed miRNAs between high and low *π*_*T*_-miR LumA-BCa samples. The x-axis represents the log_2_ fold change, and the y-axis shows the −log_10_ adjusted p-value. Vertical dotted lines indicate thresholds for |log_2_ fold change| > 1, and the horizontal line marks an adjusted p-value < 0.05 significance cutoff. (g) Bar plot summarizing significant GSEA results using genes differentially expressed between the high and low *π*_*T*_-miR groups. Hallmark pathways are ordered by normalized enrichment score, with only pathways with adjusted p-values less than 0.05 shown. For all plots, significance levels are denoted by: *, p < 0.05; **, p < 0.01; ***, p < 0.001;

To understand the shifts in the miRNA landscape between high and low *π*_*T*_-miR samples, we performed differential expression analysis of miRNAs between these groups (**Methods**). Among the top 10 most differentially expressed of these miRNAs (**Figure 4f**), miR-411-5p, miR-410-3p, miR-381-3p and miR-99a-3p have been reported to have tumor suppressive function in breast cancer^45–48^, while miR-1254 has been reported to act as a tumor suppressor by regulating HER2/HER3 signaling and can restore sensitivity to tamoxifen in HER2-positive cells^49,50^. Additionally, miR-758-3p has been profiled as part of the DANCR-miR-758-3p-PAX6 molecular network to regulate apoptosis and autophagy^51^, and miR-3609 sensitizes breast cancer cells to adriamycin chemotherapy by blocking the PD-L1 immune checkpoint^52^. The remaining miRNAs (miR-136-3p, miR-135a-5p, and miR-452-5p) functions have not been characterized in breast cancer. The identified directions of these miRNAs in association with high or low *π*_*T*_-miR do not always recapitulate their reported individual roles. This suggests that *π*_*T*_-miR may capture both known and previously unexplored concerted efforts across all miRNAs in breast cancer.

Because of the regulatory role of miRNAs on messenger RNA (mRNA)^53^, we further compared pathway-level mRNA expression profiles between high and low *π*_*T*_-miR tumors using 405 LumA samples with matched mRNA-seq data. Gene set enrichment analysis (GSEA) revealed widespread transcriptional differences across diverse hallmark pathways, encompassing proliferative, metabolic, immune, mesenchymal and stromal processes (23 pathways had |normalized enrichment scores| > 1 and adjusted p-values < 0.05, **Figure 4g, Methods**). Rather than implicating a single pathway or cellular function, these broad and multifaceted shifts suggest that variation in *π*_*T*_-miR reflects a global reprogramming of gene regulation consistent with the broad influence of miRNAs on multiple targets^54–56^. Together, these observations indicate that *π*_*T*_-miR captures a novel composite molecular phenotype linked to diverse biological processes within LumA breast cancer.

### Application to spatial transcriptomic data

Current spatial transcriptomics deconvolution methods primarily estimate cell-count proportions within tissue spots using single-cell RNA-seq references^31,57,58^. While proven valuable for characterizing tissue architecture and cellular neighborhoods^59,60^, these approaches quantify cellular presence rather than transcription activity and thus do not capture heterogeneous transcriptional states within the TME, a distinction that is critical in cancer^7,8,61^. We hypothesize that the spot-level tumor cell transcript proportion may provide complementary information beyond cell counts. To evaluate this and assess the utility of DeMixNB, we analyzed two lung cancer samples spanning distinct histologies and stages -lung adenocarcinoma (LUAD) stage III, and lung squamous cell carcinoma (LUSC). Our analytical pipeline begins with H&E slide processing using an established deep learning framework (**Methods**). This approach generates a cell-type count table for all spots, allowing us to catalog spots containing no tumor cells as reference and the rest as mixed tumor spots to deconvolve. After preprocessing (**Methods**), spot-level expression profiles for these two types of spots were used as the input data of DeMixNB (**Figure 5a**). Hence, DeMixNB does not require additional single-cell RNA sequencing (scRNA-seq) data to build reference data. DeMixNB analysis yielded tumor cell transcript proportions (*π*_*T*_), which we compared with H&E-derived tumor cell count proportions (*ρ*_*T*_) to assess the relationship between transcript activity and cellular proportions across 3,755 spots from the 2 samples (**Methods**). We found H&E-derived *ρ*_*T*_, which also does not require additional data, to be more robust than cell-type proportions estimated using public scRNA-seq as reference from RCTD^34^ and CARD^35^ (**Methods, Supplementary Figures S5-6**).

**Figure 5.**
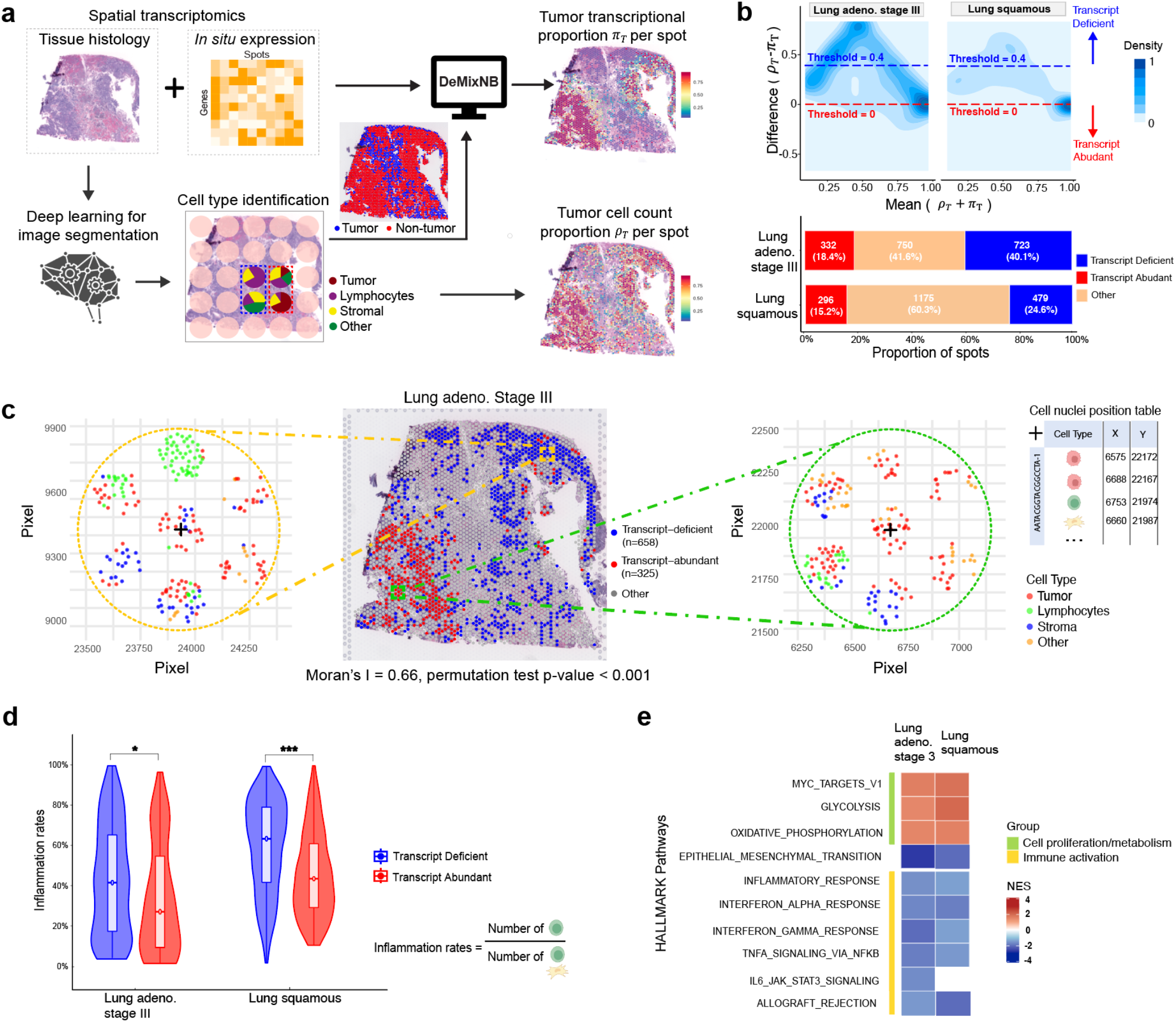
Deconvolution of spatial transcriptomics data for lung cancer. (a) Schematic illustration of the DeMixNB analytical pipeline for spatial transcriptomics data. H&E histology images undergo deep learning-based cell identification to identify tumor and multiple categories of non-tumor spots, including stromal cells and lymphocytes. Spot-level gene expression profiles from tumor-containing (mixed) spots and tumor-free (reference) spots serve as input for DeMixNB to estimate tumor transcript proportions (*π*_*T*_). H&E-derived cell identification was also used to calculate tumor cell count proportions (*ρ*_*T*_) for comparison. (b) MA density plots showing the distribution of differences (*D* = *π*_*T*_ − *ρ*_*T*_) between tumor transcript proportions and tumor cell count proportions versus their means for a lung adenocarcinoma stage III sample (left) and a lung squamous cell carcinoma sample (right). Horizontal dashed lines indicate thresholds for categorizing transcript-deficient (*D* ≥ 0.4, blue) and transcript-abundant (*D* ≤ 0; red) spots. Bar plots show the proportion of spots in each category for both samples. (c) Spatial visualization of transcript-deficient (blue) and transcript-abundant (red) spots overlaid on H&E histology image for the lung adenocarcinoma stage III sample. Middle panel shows Moran’s I spatial autocorrelation analysis demonstrating significant clustering (I=0.66, p<0.001). Representative regions (left and right) highlight spots with comparable tumor cell proportions but differential transcript activities based on local immune environment. Coordinate information is used for spatial reference. (d) Comparison of local inflammation rates between transcript-deficient and transcript-abundant spots across both lung cancer samples. Inflammation rates represent the proportion of lymphocytes relative to the total lymphocyte and stromal cell count for each spot and its six adjacent neighbors. Statistical significance was assessed using the Wilcoxon rank-sum test with Benjamini-Hochberg (BH) correction. (e) Gene set enrichment analysis (GSEA) results showing significantly enriched Hallmark pathways in transcript-deficient versus transcript-abundant spots. Pathways are ordered by normalized enrichment score (NES), with only pathways meeting significance thresholds (BH-adjusted p < 0.05) displayed. For all plots, significance levels are denoted by: *, p < 0.05; **, p < 0.01; ***, p < 0.001.

To assess whether the differences between *π*_*T*_ and *ρ*_*T*_ reflect meaningful biological variation, we measured the spot-level difference (D) as *ρ*_*T*_ − *π*_*T*_, and used the difference to identify three groups of spots (**Figure 5b, Methods**): transcript-deficient spots where tumor cells maintain physical presence but exhibit substantially reduced transcriptional activity (D > 0.4), transcript abundant spots where transcript proportions meet or exceed cellular proportions (D < 0), and the rest. In the LUAD Stage III and LUSC samples, the transcript-deficient spots comprised 40.1% and 24.5% of spots, and the transcript-abundant spots comprised 18.4% and 15.2% of spots, respectively, demonstrating marked spatial variation in tumor transcriptional activity in both samples. Using Moran’s I, which measures whether different groups of spots cluster together more than expected by chance (0=random, 1=clustered, −1=dispersed, **Methods**), we found both samples showed strong positive spatial clustering, with Moran’s I values of 0.66 (**Figure 5c**) for LUAD stage III and 0.59 (**Supplementary Figure S7**) for LUSC (permutation test p < 0.001 for both, **Methods**). This non-random spatial clustering suggests potential spatial biological factors that may regulate tumor cell activities. Detailed examination of representative regions revealed that spots with comparable tumor cell proportions exhibited markedly different transcript activities depending on their local immune environment: a higher number of immune cells is observed in the neighborhood of transcript-deficient spots (see an example neighborhood in **Figure 5c**). While recent multimodal spatial-omics study^62^ has characterized the co-evolution of epithelial alveolar progenitors and proinflammatory niches in lung adenocarcinoma, the differential transcriptional response of tumor cells to this ecosystem remains to be fully quantified. We therefore further explored the relationship between immune infiltration and tumor transcript suppression by calculating a local inflammation rate across all spots. For each spot defined as transcript-deficient or abundant by *ρ*_*T*_ − *π*_*T*_, we summed lymphocytes counts across the spot itself and its six adjacent neighbors, divided by the total lymphocytes and stroma cells in the same region to obtain a measure of local inflammation rate (**Figure 5c, Methods**). Transcript-deficient spots consistently showed significantly higher inflammation rates compared to transcript-enriched spots across both samples (adjusted p-values < 0.05, **Figure 5d**). This inverse relationship suggests that local immune pressure may actively suppress tumor cell transcript production, possibly by driving malignant cells into quiescent KRT8^+^ alveolar intermediary cells (KAC) state as proposed in recent literature^63–65^. A change of cutoff value on *ρ*_*T*_ − *π*_*T*_ from 0.4 to 0.3 to include more transcript deficient spots replicated the above observations in both samples (**Supplementary Figure S8**), supporting robustness of our findings. For the LUSC sample where we did not find strong evidence to rule out CARD-based *ρ*_*T*_, we also identified transcript-deficient and transcript-abundant populations using this alternative set of tumor cell count proportion estimates and found a consistent inverse relationship between immune infiltration and transcriptional activity (**Supplementary Figures S9, Methods**).

To validate the biological processes underlying transcript deficiency, we performed GSEA on differentially expressed genes between transcript-deficient and transcript-abundant spots in both samples (**Methods**). The identified pathways were highly consistent with the recently discovered inflammatory-KAC axis through multi-omic and high-resolution spatial profiling of the same sample^62^. The most prominently enriched pathways in transcript-deficient spots point to immune activations while those enriched in transcript-abundant spots correspond to cell proliferation and metabolism (**Figure 5e**). Crucially, Epithelial-Mesenchymal transition (EMT) was also significantly enriched in these transcript-deficient spots. The co-occurrence of high inflammation, tumor cell transcript reduction, and EMT signatures suggests that immune infiltration may induce tumor lineage plasticity, forcing malignant cells to dedifferentiate into a stress-resistant, low-transcript phenotype to survive the inflammatory microenvironment^4,66,67^. These findings demonstrate that DeMixNB-derived *π*_*T*_ captures functionally distinct tumor cell transcriptional states which are clustered across spatial regions within a tumor sample, providing an additional aspect in understanding the tumor heterogeneity in the complex TME.

## Discussion

We develop DeMixNB to address a methodological gap by enabling accurate deconvolution of sparse count sequencing data through a semi-reference-based approach with negative binomial distributions^12^. While existing methods are primarily designed for cell-type count deconvolution, DeMixNB specifically targets tumor transcript proportion estimation and accommodates the overdispersion characteristic in miRNA-seq and spatial transcriptomics data. It is important to measure tumor transcript proportion for cancers because it provides the foundation to evaluate features of tumor cell plasticity^61^. We also generate a unique benchmark sparse-count mixed cell-line dataset with known cell count and transcript proportions to demonstrate the good performances of DeMixNB. Only requiring adjacent mixed non-tumor reference data rather than single-cell derived reference profiles, which are often unavailable or technically challenging to produce, DeMixNB provides a practical solution for quantifying tumor transcript proportions (*π*_*T*_) in real-world sequencing contexts. The DeMixNB algorithm has been implemented as a function in the R package DeMixT2.0, available from the GitHub repository: https://github.com/wwylab/DeMixT/tree/develop, and will be made available through Bioconductor at https://www.bioconductor.org/packages/release/bioc/html/DeMixT.html.

By deconvolving sparse transcripts using DeMixNB, we are able to uncover novel biological nuance when tumor cell purity (*ρ*_*T*_) and tumor transcript proportion (*π*_*T*_) diverge. In spatial transcriptomics data, *π*_*T*_ helps identify tumor transcript-deficient spots, where we observe non-random spatial clustering of these spots near immune-infiltrated regions, with these areas showing significantly elevated lymphocyte proportions. Although spatial immune-tumor interactions have been studied in recent spatial transcriptomic literature^62,68–70^, DeMixNB quantifies the responsive tumor-cell specific overall transcriptional activity for the first time. Similarly, the novel prognostic utility of *π*_*T*_-miR independent of DNA copy-number based purity estimates in TCGA breast cancer further validates this functional measure. In this scenario, elevated tumor miRNA proportions may reflect upregulation of oncogenic miRNAs or enhanced regulatory plasticity enabling cancer cell adaptability under selective pressures. Tumor total miRNA proportions represent a previously unexplored feature of cancer, and may lead to more refined understanding of the regulatory transformations of malignancy.

Future application across diverse cancer types may reveal pan-cancer trends to establish clinically robust biomarkers based on tumor transcript proportions for sparse-count RNA species. By adjusting further for tumor purity and genome dosage effects, a more accurate measure of total tumor miRNA content per tumor cell could be calculated, which may serve as a novel biomarker^12,61^. The spatial transcriptomics capability opens critical avenues for developing region-specific signatures that can deconstruct complex bulk transcriptomic signals and quantify true tumor-intrinsic transcriptional activities. A key translational objective involves determining whether baseline proportions of transcript-deficient spots predict clinical trajectories such as metastatic progression or immunotherapy response, given their enrichment for immune pathways and potential role in checkpoint blockade outcomes.

Most existing deconvolution benchmarking approaches rely on the assumption that transcript proportions directly correspond to cellular proportions, which may not fully capture cell-type-specific differences in transcriptional output and RNA content^12^. To provide a more comprehensive evaluation framework, we developed an experimental benchmarking approach that establishes ground truth based on actual transcript proportions rather than cellular ratios. By measuring miRNA density per cell and mixing extracted miRNAs rather than cells themselves, we generated samples with precisely known transcript proportions. Furthermore, the inclusion of both *Dicer1* wild-type and knockout cell mixtures simulates populations with unknown tumor subtypes, reflecting the heterogeneous nature of real-world clinical samples where distinct molecular subtypes are typically uncharacterized prior to deconvolution. This approach provides a biologically grounded benchmarking standard and represents a valuable community resource for evaluating deconvolution methods across realistic scenarios.

Although DeMixNB has shown its strength in transcript proportion deconvolution of sparse count RNA-seq, its accuracy by using a discrete count distribution is achieved by sacrificing computational speed, as compared to a continuous normal or log-normal distribution. When the count data is not sparse, DeMixNB loses more on speed than gaining accuracy as compared to DeMixT. In addition, we observe some numerical challenges during estimation due to the nature of semi-reference-based inference. We mitigate the numerical instability by running the algorithm multiple times with different starting values and generate a consensus as the final estimates, further increases the computing time. Optimizations can be made to speed up, e.g., adding more parallelization, or implementing the likelihood calculation in Python to utilize GPUs. Using representative reference data is essential for running any semi-reference-based deconvolution methods including DeMixNB. Finally, our digital pathology-derived cell count proportions were limited to four cell types, so that important immune cell populations, such as macrophages, could not be determined. This limitation restricted our current analysis from gaining deeper insights into the interactions between tumor transcripts and immune responses. Future work may incorporate more comprehensive cell type annotations using advanced digital pathology tools or complementary technologies such as multiplexed immunofluorescence. This would enable a more detailed characterization of the tumor microenvironment and a better understanding of its impact on tumor cell transcriptional activities.

## Conclusion

Modeling sparse count data as those in spatial transcriptomics and miRNA-seq to evaluate tumor cell plasticity presents unique challenges. We developed a method, DeMixNB, that can deconvolve the tumor transcript proportions in such data without the need for single-cell-level reference data. Using a novel benchmarking experimental dataset, DeMixNB demonstrated better performances than existing methods. When applied to real data, DeMixNB estimates of transcript proportions capture distinct biological information not otherwise reflected in tumor purity estimates. The framework established by DeMixNB extends beyond mRNA and miRNA applications to encompass emerging bulk sequencing modalities profiling the broader RNA species, including piRNAs, snoRNAs, and lncRNAs. Therefore, DeMixNB as a tool will empower future studies interrogating the complex biology of cancer RNomes.

## Method

### Datasets representing sparse transcriptomes for deconvolution

#### Experimental mixed miRNA-seq data with known mixing proportions

To generate a benchmarking dataset with known ground truth proportions, HCT116 colon cancer cells and HS-5 stromal cells were cultured at 37°C in a humidified incubator with 5% CO_2_. HS-5 cells were maintained in Dulbecco’s Modified Eagle’s Medium (DMEM;Corning) supplemented with 10% fetal bovine serum (FBS;Corning) and 1% Antibiotic-Antimycotic (Gibco), while HCT116 cells were maintained in McCoy’s 5A medium (Corning) with the same supplements. The Dicer1 knockout (KO) HCT116 cells were cultured under the same conditions with the addition of G-418 selective antibiotic solution (neomycin resistance) to preserve the homogenous KO population. Total RNA was extracted from 1 x 10^6^ cells using the Zymo RNA Isolation Mini Kit (Zymo Research). Small RNA concentrations (ng/µL) were quantified using a Qubit 4 Fluorometer (Invitrogen) with the Qubit microRNA Assay Kit (Invitrogen). Based on these measurements, RNAs from each cell type were mixed in defined proportions to generate samples reflecting predetermined tumor:stromal total miRNA ratios: 0:100, 30:70, 50:50, 60:40, 70:30, 85:15, and 100:0. Each set of mixtures was prepared independently for parental HCT116 combined with HS-5 RNA and Dicer1 KO HCT116, combined with HS-5 RNA with three biological replicates per condition, resulting in 39 total samples. Sample quality was validated using an Agilent Bioanalyzer (Agilent). miRNA sequencing libraries were prepared using the QIAseq miRNA Library Kit (Qiagen) and sequenced on an Illumina platform by SeqMatic LLC.

Raw sequencing data quality was assessed using FastQC, and adapter sequences were trimmed using Trim Galore. The processed reads were then aligned to miRBase (version 22.1) and the human reference genome GRCh38 using the miRDeep2 pipeline to quantify miRNA expression levels. This alignment produced a count matrix of mature miRNA expression for each sample, with annotations from the v22.1 miRBase release^71^. Raw files and aggregated expression matrix for these benchmarking samples have been deposited at GEO.

#### MiRNA-seq data from patients with breast cancer

Isoform level miRNA expression data for TCGA-BRCA breast cancer samples were downloaded from the Genomic Data Commons (GDC) data portal (https://portal.gdc.cancer.gov). The miRNA count matrix was upper quartile normalized, and miRNAs detected in 10 or fewer samples were removed. Additionally, miRNAs with a mean expression greater than 10,000 in either the normal reference or mixed samples were excluded to reduce computational load and ensure adherence to the assumptions of the negative binomial model.

Breast cancer intrinsic subtype prediction was performed using the R `genefu` package, based on the ‘PAM50’ subtyping signature. Transcript expression data for subtype prediction and clinical annotations (including stage and progression-free survival) were retrieved from the GDC data portal.

### Spatial transcriptomic data from patients with lung cancer

The spatial transcriptomic data for Lung adenocarcinoma stage III sample was generated using 10x Genomics’ Visium technology on formalin-fixed, paraffin-embedded (FFPE) tissues. RNA was extracted with the Qiagen RNeasy FFPE Kit and quality-checked using the Agilent RNA 6000 Pico Kit (DV200 > 50%). Tissue sections (6.5×6.5 *mm*) underwent deparaffinization, H&E staining, decrosslinking, probe hybridization, and library construction, using the Visium Spatial Gene Expression Slide. The Visium spatial gene expression FFPE libraries were constructed using the Visium CytAssist Spatial Gene Expression for FFPE Human Transcriptome Probe Kit (PN-1000444) following the manufacturer’s guidance.

The spatial transcriptomic data for lung squamous sample were generated using 10x Genomics’ Visium CytAssist Spatial Gene Expression platform on formalin-fixed, paraffin-embedded tissues (Avaden Biosciences). Tissue sections (5 µm) were deparaffinized, H&E-stained, decrosslinked, and processed for probe hybridization and library construction following the manufacturer’s demonstrated protocols (CG000518, CG000520). Visium FFPE libraries were constructed using the Visium CytAssist Spatial Gene Expression for FFPE Reagent Kit (CG000495) according to the standard workflow. The Lung squamous spatial transcriptomic dataset was obtained from the 10x Genomics public data repository:http://www.10xgenomics.com/datasets/human-lung-cancer-ffpe-2-standard.

### The DeMixNB model

Due to the lack of tumor reference profiles, we develop the DeMixNB model to be semi-reference-free, requiring only nontumor expression data at the same level as the mixed expression data. Similar to previous deconvolution methods in the literature, we adopt the negative binomial distribution at the core of our DeMixNB model to address the sparse count issue present in the miRNA-seq data and the spatial transcriptomics data (**Supplementary Figures S1-2**). We focus on the deconvolution of transcript proportions in the case of two cell compartments, tumor cells and non-tumor cells, only. Denote the observed mixed expression at sample *i* for gene *g* as *Y*_*ig*_. The deconvolution model assumes that it can be decomposed into individual contributions of each cell type, i.e. *Y*_*ig*_ = *T*_*ig*_ + *N*_*ig*_, where *T*_*ig*_ is the contribution of the tumor component and *N*_*ig*_ is the contribution of the non-tumor component at sample *j* for gene *g*. Both *T*_*ig*_ and *N*_*ig*_ are not observed.

We assumed that both *T*_*ig*_ and *N*_*ig*_ follow a negative binomial distribution parameterized as *T*_*ig*_ ~ *NB*(*π*_*i*_*µ*_*tg*_, *ϕ*_*tg*_) and *N*_*ig*_ ~ *NB* /(1 − *π*_*i*_)*µ*_*ng*_, *ϕ*_*ng*_4, where *π*_*i*_ represents the proportion between the tumor and non-tumor components, *µ*_*tg*_ and *µ*_*ng*_ are the means of the distributions, and *ϕ*_*tg*_ and *ϕ*_*ng*_ are the dispersion parameters. Assuming independence between *T*_*ig*_ and *N*_*ig*_, the expected expression of *Y*_*ig*_ can be expressed as *E*(*Y*_*ig*_) = *E*(*T*_*ig*_) + *E*(*N*_*ig*_) = *π*_*i*_ *µ*_*tg*_ + (1 − *π*_*i*_) *µ*_*ng*_. This is a variation that approximates the setup in previous studies^12,13^, which state *Y*_*ig*_ = *π*_*i*_ *T*_*ig*_ + (1 − *π*_*i*_) *N*_*ig*_. Therefore, a major assumption introduced by DeMixNB is the proportions only affect the mean expressions and do not scale the dispersion levels across genes. This assumption deviates from real scenario but in return gains computational tractability of model parameters. In theory, if one has access to the true expressions of *T*_*ig*_ and *N*_*ig*_, as is the case in a simulation setting, the true transcript proportion can then be calculated as 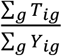. However, in practice, we estimate an approximation of the true transcript proportion using ***π***, which has been previously demonstrated for the model *Y*_*ig*_ = *π*_*i*_ *T*_*ig*_ + (1 − *π*_*i*_) *N*_*ig*_^61^.

Based on the DeMixNB model detailed above, we implemented the DeMixNB algorithm to estimate the parameters *π*_*i*_, *µ*_*tg*_, *ϕ*_*tg*_, *µ*_*ng*_, and *ϕ*_*ng*_ that maximizes the total likelihood (see a pseudo-code description of the algorithm in the **Supplementary Information**). The input data for DeMixNB includes two data matrices: 1. ***Y***, the matrix containing the mixed expression data, which is the target of deconvolution; 2. ***Y***^ref^, the matrix containing reference data of only non-tumor expressions. The output of DeMixNB would be the estimated transcript proportions in the mixed data ***Y***, as well as the other parameter estimates. We require that samples of reference data for the non-tumor component be provided, i.e. samples where *π*_*i*_ = 0 for all *i*, to estimate the parameters *µ*_*ng*_ and *ϕ*_*ng*_ first. Therefore, our method falls into the category of semi-reference-based deconvolution. To estimate the remaining parameters, we take the estimated distributions of the non-tumor components as fixed, then perform ICM^36^. ICM has been proven to guarantee convergence to local maximum, a desirable property for parameter estimation. The ICM algorithm iteratively updates the mixing proportions ***π*** and distribution parameters ***μ***_***t***_ and ***ϕ***_***t***_, maximizing the conditional likelihoods at each iteration. This process is similar to the DeMixT algorithm proposed in Wang et al^12^.

The total likelihood for observing the matrix *Y* can be expressed as *L*(***Y***) = ∑_*i*_ ∑_g_ *L*(*Y*_*ig*_), assuming the expression level for each gene *g* at each sample *i* is independent from one another. The individual likelihood of *Y*_*ig*_ can be calculated by a convolution of the two individual probability distributions of *T*_*ig*_ and *N*_*ig*_.

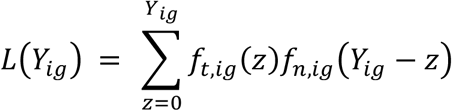

where *z* is the iterating variable and

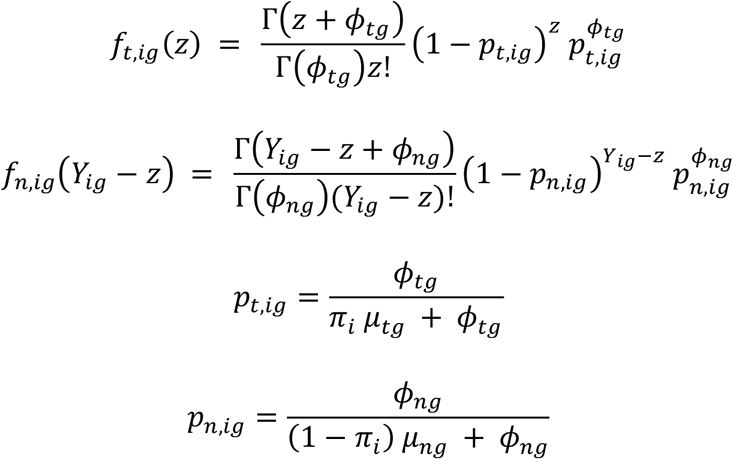

#### Numerical stability and software

The DeMixNB algorithm minimizes the negative log-likelihood instead of the likelihood for improved stability. Due to the high-dimensional search space and the ICM’s converging to local minimum, we found that the initiating proportion values can result in final estimates hitting the optimizer bounds incorrectly on rare occasions. Therefore, by default, our DeMixNB function will start the algorithm three times with different seeds (randomly generated starting values) and perform a consistency check across the runs. At each spot or sample, we take the maximum difference between the three estimates and if the maximum difference is less than 0.1, we deem the estimates for the spot as consistent and reliable. The issue of converging at distinct values across three runs is rarely seen in simulations. When it does occur, using the average estimated proportion across consistent samples provides a reasonable solution. We also set the bounds of the proportion estimates to be 0.05 and 0.95. The DeMixNB software is mainly based on the R programming language, e.g. using the built-in L-BFGS-B optimizer, with some computation-heavy components implemented in the C programming language with parallel computing support. The DeMixNB() and its documentation has been integrated in the DeMixT^24^ R package v2.0.

### Model validation using simulated sparse count data

#### Study setup

To evaluate the performance of DeMixNB, we considered three simulated scenarios, each generating simulated data based on negative binomial distributions, with varying prior distributions for the mean and dispersion parameter. To generate the expression matrices, we first sample the true proportions *π*_*i*_ from a uniform distribution and the mean and dispersion parameters *µ*_*tg*_, *µ*_*ng*_,*ϕ*_*tg*_, *ϕ*_*ng*_ from predefined prior distributions, detailed in the **Simulation Settings** section. The mixed expression counts are subsequently generated as *Y*_*ig*_ = *T*_*ig*_ + *N*_*ig*_ = *NB*(*π*_*i*_*µ*_*tg*_, *ϕ*_*tg*_) + *NB* /(1 − *π*_*i*_)*µ*_*ng*_, *ϕ*_*tg*_4. In each scenario, 100 datasets are generated with the exact same true proportions *π*, and the expressions are sampled from the same negative binomial distributions with different seeds. The true transcript proportions based on the generated data, calculated as 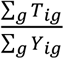, are recorded for evaluation. With the generated data as input, we run the DeMixNB function with default settings to perform deconvolution. The estimated proportion vector is output by the algorithm for each dataset, denoted as 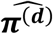 for dataset *d*, which is compared with the true proportion vector. We also compared DeMixNB to three state-of-the-art partial-reference deconvolution methods: DeMixT^15^, ISOpure^16^, and PREDE^19^. DeMixT (version 1.20.0) is capable of three-component deconvolution, but we are only using its two-component deconvolution functionality. Additionally, we did not use its built-in preprocessing and gene selection methods to ensure a fair comparison (by setting `if.filter` to `FALSE`). As DeMixNB is built as a variant of DeMixT, DeMixT also requires a set of reference samples in order to estimate the distribution parameters of the non-tumor component first. For ISOpure (version 1.1.3), we only use its first step, the `ISOpure.step1.CPE` function with default parameters, and extract the `alphapurities` from its output as the estimated proportions. PREDE (version 1.2.1) requires the reference of the known expression profiles exactly. In this case, we treat the non-tumor component as one cell type and provide the empirical means of the generated expressions as the reference expression profile. We set type=“GE”, K = 2 and `iters=1000` in its function parameter to enforce two-component deconvolution, using the average expression profiles of the normal samples as reference.

#### Simulation settings

In all simulations, the sample size for the mixed expression matrix is set to 100 and that for the normal reference matrix is set to 50. The number of genes is fixed at 1000. And the prior distributions for the dispersion parameters *ϕ*_*tg*_ and *ϕ*_*ng*_ are set to while we focus on the impact of the mean parameter:

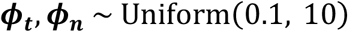

We consider three different simulation scenarios with varying prior distributions for *µ*_*tg*_, *µ*_*ng*_. In scenario A, we mimic a non-sparse dataset by setting high values *µ*_*tg*_ and *µ*_*ng*_ with means around 200:

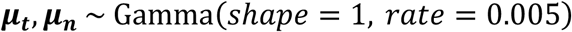

In scenario B, we make the data sparse by reducing the means for *µ*_*tg*_ and *µ*_*ng*_ to around 20:

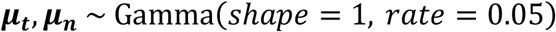

In scenario C, we consider the case where genes might express based on different distributions for the same cell type. Specifically, we consider two distributions for the tumor cells and assign 800 genes to the first distribution and 200 genes to the second. Both distributions have small means to ensure the final mixed expression matrix remains sparse.

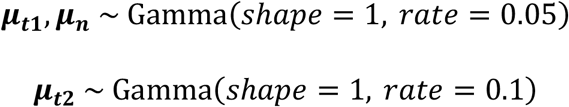

To evaluate the performance of the model, we calculate the average absolute bias across the 100 estimates, the average root mean squared error (RMSE), and the average concordance correlation coefficient (CCC) between the estimated and the true proportions. For each dataset, we also calculate a consistency metric (sumEst) where we use the parameters for the tumor component to generate reference data to estimate the proportion of the non-tumor component and verify that the sums of the estimates for the non-tumor component and those for the tumor component are close to 1.

### Model validation using experiment data as benchmark

#### Experimental setup

In our benchmarking dataset, we have reference profiles for the pure fibroblasts, *Dicer1* wildtype cancer cells, and *Dicer1* knockout cells. We benchmarked deconvolution performance for all methods using all samples mixed from both the wildtype and knockout cells, i.e., the tumor-specific total miRNA transcripts vary largely across samples, as this represents a realistic biological scenario, where cancers in a cohort may come from distinct subtypes with different expression profiles. We evaluated DeMixNB against three established semi-reference-based deconvolution methods, DeMixT, ISOpure, and PREDE, as well as a reference-based deconvolution method for miRNA-seq, DeconmiR. Prior to deconvolution analysis, the miRNA count matrix was normalized using upper quantile normalization. The normalized expression matrix was then used as input for all methods.

#### Deconvolution of benchmarking data

For DeMixNB, pure HS-5 fibroblast samples (MIX-1-1, MIX-2-1, MIX-3-1) were used as the reference normal component. The remaining mixed samples were treated as the input bulk tissue samples requiring deconvolution. DeMixT version 1.20.0 was used with parameters set to `gene.selection.method=“GS”`, `ngene.Profile.selected=200`, `ngene.selected.for.pi=200,niter=65`,`nbin=50`,and `filter.sd=2`, and used the count matrices of the pure fibroblast samples as normal reference. For PREDE (version 1.2.1), we implemented the PREDE function with parameters `type=“GE”`, `K=2`(representing two cell types), and `iters=1000`, using the average expression profile of the pure HS-5 samples as the reference normal component (W1). For ISOpure (version 1.1.3), the `ISOpure.step1.CPE` function from the ISOpureR package^37^ was run using default parameters. For DeconmiR^33^ (version 0.0.0.9000) since the model requires a full set of reference profiles for both T and N, we set up the T component in three ways: 1) the average expression of *Dicer1* wildtype and knockout cell lines for all mixed samples; 2) the average expression of one type of tumor cells: wildtype for all, knockout for all; 3) the average expression of matched tumor cell types: wildtype for wildtype, knockout for knockout. These settings deviate further and further away from real data scenarios for deconvolution. Deconvolution was performed using (`method=“RPC”`).

Model performance was evaluated by calculating the root mean square error (RMSE) and concordance correlation coefficient (CCC) between the ground truth mixing proportions and the estimated tumor proportions from each method. SumEST was not used in this evaluation, as a single reference profile for the both cancer cell populations is not feasibly obtained. We used bootstrapping to measure uncertainty. From the mixed samples, we randomly sampled sets of 36 samples from the mixed-cell benchmarking dataset, 100 times with replacement. If a sampling didn’t include 3 different mixing proportions, this sample would be redrawn. The Wilcoxon rank-sum test was used to assess the differences between paired CCC and RMSE distributions from the above methods.

### Analysis of TCGA-BCa miRNA-seq data

#### Deconvolution

Deconvolution of tumor miRNA proportions was carried out for 899 TCGA breast cancer samples, using matched normal tissues from 98 samples as normal reference. Prior to deconvolution, input count matrices were upper-quartile normalized, and miRNAs with mean expression across samples > 10,000 were filtered out, resulting in a set of 1,191 miRNAs. DeMixNB was run with the scaling factor set to 0.1, which multiplies input count data by a factor of 0.1, which improves reliability of parameter estimation for highly expressed miRNAs and improves the runtime. Following deconvolution, samples were filtered to those with clinical annotations including stage and PAM50 subtype.

#### Comparison with tumor purity estimates

ASCAT-based tumor purity values were obtained from the Van Loo Lab GitHub repository (https://github.com/VanLoo-lab/ascat/tree/master/ReleasedData). Pearson correlation was calculated between tumor purity values and *π*_*T*_-miR estimates.

#### Survival analysis

Within the Luminal A subgroup, tumor purity–associated differences in progression-free survival were evaluated using recursive partitioning, Kaplan–Meier analysis with log-rank tests, and Cox regression. Recursive partitioning was performed with the rpart package (v4.1.23) to identify an optimal cutoff for *π*_*T*_-miR and ASCAT-based tumor purity using parameters that restricted the tree to a single split (maximum depth = 1), required a minimum of five observations to attempt a split, a minimum terminal node size of 10% of the samples, and a complexity parameter set to a low value (cp = −0.01) to allow the model to identify a single, unbiased cutoff point without pruning additional splits. Progression-free survival differences between the resulting high *π*_*T*_-miR and low *π*_*T*_-miR groups, along with high and low ASCAT-derived purity, were assessed using Kaplan–Meier analysis with log-rank tests. A multivariate Cox proportional hazards model incorporating age and the clinical pathological stage was also used to evaluate the independent prognostic value of *π*_*T*_-miR.

#### Differential expression and gene set enrichment analysis

Differential expression analysis of miRNA and mRNA transcripts was performed using `DESeq2` (v1.44.0). Gene set enrichment analysis (GSEA) of differentially expressed genes was conducted using `fgsea` (1.30.0) with genes ranked by normalized enrichment scores (NES) from DESeq2. Hallmark pathway gene sets were obtained from the Molecular Signatures Database (http://www.gsea-msigdb.org/gsea/msigdb/collections.jsp).

### Deconvolution of spatial transcriptomic data

#### Deep learning-based cell classification

Cell detection and classification are completed using a deep learning pipeline ^72,73^. Initially, the whole-slide H&E images were segmented using a Micro-Net^74^ architecture to remove background noise and artifacts. Then the nuclei within the segmented tissue regions were identified through a Spatially Constrained Convolutional Neural Network (SCCNN) model for cell detection. Cell classification is performed using a neighboring ensemble predictor and voting strategy. Cells in the tissue are categorized into four groups: (malignant epithelial cells), lymphocytes (including plasma cells), non-inflammatory stromal cells (fibroblasts and endothelial cells), and another cell type that includes non-identifiable, and less abundant cells such as macrophages.

#### Data pre-processing

Raw spatial transcriptomic data were preprocessed to ensure data quality and suitability for downstream analyses. Initial gene filtering removed mitochondrial genes (MT-), microRNA genes (MIR-), long intergenic non-coding RNAs (LINC), immunoglobulin genes (IG*), and T-cell receptor genes (TR*). Genes detected in fewer than 10 spatial spots were excluded. Subsequently, spatial spots were filtered based on minimum library size (≥600 total unique molecular identifiers, UMIs) and gene detection thresholds (≥300 detected genes). H&E-based cell-type proportions (tumor, lymphocytes and stroma) and cell counts per spot are summarized and used to identify tumor-mixed and non-tumor spots. Spots containing fewer than five total cells were excluded. Spots without tumor cells identified through this pipeline served as reference spots for subsequent tumor-specific deconvolution. After filtering, we have 2,252 mixed spots and 834 reference spots for the LUAD stage III sample and 2,457 mixed spots and 1,214 reference spots for the LUSC sample.

#### Spot grouping

We selected thresholds for classifying spots as tumor transcript-deficient or tumor transcript-abundant using a Bland-Altman representation of digital pathology-derived tumor cell count proportion (*ρ*_*T*_) and DeMixNB-derived tumor cell transcript proportion (*π*_*T*_). To ensure robust analysis, we restricted our selection to spots with at least moderate tumor content (*ρ*_*T*_ ≥ 0.25), as difference *D* in regions with very low tumor purity are difficult to interpret biologically and may reflect measurement noise rather than real transcriptional differences. We defined transcript-deficient spots as those with *D* ≥ 0.4, or 0.3, where *ρ*_*T*_ substantially exceeded *π*_*T*_, and transcript-abundant spots as those with *D* ≤ 0, where *π*_*T*_ met or exceeded *ρ*_*T*_. These thresholds were chosen based on the observed density distribution patterns to capture biologically distinct populations while maintaining sufficient sample sizes for downstream statistical analysis.

#### Gene selection for deconvolution

While gene selection is not necessary for the accurate deconvolution using DeMixNB, reducing the gene size can substantially decrease the computing time. For spatial transcriptomics technologies such as the 10x Visium, they sequence approximately 18,000 genes, which is too many to add up for a count-based model. We first normalized and scaled the spatial transcriptomics data using the `SCTransform` function from the `Seurat` R package (v4.0). Principal component analysis (PCA) was performed on the resulting SCT assay data with the `RunPCA()` function. Graph-based clustering `FindNeighbors()` and `FindClusters()`, resolution = 0.8) and uniform manifold approximation and projection (UMAP; `RunUMAP()`) embedding were subsequently performed using the first 50 principal components. Spatially variable genes were then identified using the Moran’s I statistic (`FindSpatiallyVariableFeatures(), selection method = “moransi”`), computed on all genes identified by SCTransform. Genes were ranked according to observed Moran’s I values, and the top-ranked genes (typically top 1,000 genes) were selected for deconvolution.

#### Reference-based deconvolution

We compared the H&E-based tumor cell count proportions with widely used single-cell reference-based methods (e.g., RCTD^34^, CARD^35^) derived proportions to evaluate consistency and robustness. The raw count matrices were preprocessed using the same preprocessing pipeline as for DeMixNB. Single-cell reference datasets were obtained from previous studies: one lung adenocarcinoma dataset containing 20 samples^75^ and one lung squamous dataset containing 8 samples^76^. Both reference datasets were preprocessed to retain only well-annotated cells, and each cell type was limited to a maximum of 1,000 cells to prevent bias from dominant cell types. For CARD analysis, we followed the standard tutorial on GitHub to create CARD objects with default parameters. For RCTD analysis, we created spatialRNA objects with `max_cores=4`, `doublet_mode=‘full’`. For both methods, the spot-level malignant cell proportion served as the tumor cell count proportion *ρ*_*T*_ in the downstream analysis.

#### Spatial clustering analysis and permutation test

To assess spatial clustering of transcript-deficient and transcript-abundant spots, we computed Moran’s I on the binary group label using row-standardized spatial weights (`nb2listw, style = “W”, zero.policy = TRUE`) derived from an adaptive neighbor graph constructed by distance-based criteria with k-nearest-neighbor fallback. Moran’s I measure spatial autocorrelation ranging from −1 (perfect dispersion) to 1 (perfect clustering), with 0 indicating random spatial distribution. Statistical significance was assessed by a Monte Carlo permutation test (`moran.mc, nsim = 10000`). Under the null hypothesis of spatial randomness, group labels were randomly reassigned across spots while maintaining the spatial weight structure. This process was repeated 10,000 times to generate an empirical null distribution of Moran’s I values. The one-sided p-value was calculated as the proportion of permuted statistics greater than or equal to the observed value, testing for positive spatial autocorrelation. An observed Moran’s I consistently exceeding all permuted values indicates statistically significant spatial clustering of spot groups.

#### Spot-specific inflammation rate

To quantify local tumor microenvironment composition, we calculated an inflammation rate representing the abundance of lymphocytes versus stromal cells within each spatial transcriptomic spot and its immediate neighbors. We determined spatial adjacency between spots based on Euclidean distance of their coordinates, identifying neighbors within a predefined threshold (140 *µm*). For each target spot, the total stromal and lymphocytes counts were summed across the target spot and its neighboring spots, forming a localized pseudo-bulk composition. The inflammation rate was computed as

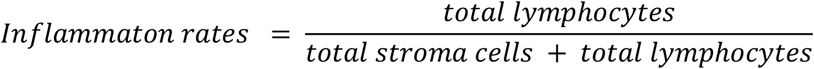

Spots with zero total stromal and lymphocytes cells were excluded from scoring to avoid division by zero. The calculated inflammation rates were then compared statistically between two defined spot groups (“transcript-deficient” and “transcript-abundant”) using the Wilcoxon rank-sum test implemented with the `wilcox.test()` function with Benjamini-Hochberg (BH) correction in R.

## Declarations

### Ethics approval and consent to participate

The study is conducted in accordance with the Declaration of Helsinki and approved by the Institutional Review Board of University of Texas, MD Anderson Cancer Center (protocol code xxx)(need to request from collaborator). All participants provided written informed consent.

### Consent for publication

All authors have approved the publication of manuscript.

### Availability of data and materials

MiRNA and mRNA expression data, along with clinical annotations for TCGA-BRCA breast cancer samples were downloaded from the Genomic Data Commons (GDC) data portal (https://portal.gdc.cancer.gov).

Lung adenocarcinoma stage III spatial transcriptomic data were deposited in the Gene Expression Omnibus (GEO) under the accession number GSE307534. The 10X Visium lung squamous cancer data is available at http://www.10xgenomics.com/resources/datasets/human-lung-cancer-ffpe-2-standard.

## Acknowledgements

M.D.M, H.Y., L.X. and W.W. was supported in part by NCI grant 1R01CA268380. P.Y. was supported in part by NCI grant 1R01CA183793. Dr. Calin is the Charles B. Barker Chair. Work in G.A.C.’s laboratory, including work from K.N. is supported by NCI grant 1R01CA222007-01A1, NIDCR grant R01DE032018, DoD CDRMP Idea Award BC200208P1 (W81XWH-21-1-0030), Team DOD grant in Gastric Cancer CA200990P2 (W81XWH-21-1-0715), DoD Idea Award PC230419 (HT9425-24-1-0052), the 2019 Faculty Achievement Award, CLL Global Research Foundation 2019 grant, CLL Global Research Foundation 2020 grant, CLL Global Research Foundation 2022 grant, CLL Global Research Foundation 2024 grant, The G. Harold & Leila Y. Mathers Foundation, two grants from Torrey Coast Foundation, an Institutional Research Grant 2024 a Development Grant associated with the Brain SPORE 2P50CA127001, an Institutional Bridge Funding grant 2023, and the Ben and Catherine Ivy Foundation grant. This work is supported by Spatial Ecology & QUantitative pathOlogy Image Analytical platform (SEQUOIA) through the MD Anderson STrategic Research Initiative Development Program (STRIDE). X.P. and Y.Y. were supported in by Lyda Hill Philanthropy FP00020550.

## Author contributions

M.D.M., H.Y., L.X., W.W. conceived the project. L.X. P.Y., and W.W. developed the mathematical method. M.D.M., H.Y., and L.X. performed the data analysis and interpretation, planned the figure design, and wrote the manuscript collaborating with all the other authors. R.L. assisted with data maintenance, manuscript writing, and figure design. M.D.M. K.N. W.W. G.A.C. conceived the benchmarking dataset design. K.N. G.A.C. generated benchmarking data. X.P. performed digital pathology of ST samples. C.E. reviewed pathology annotation and provided interpretation and domain expertise. A.S., T.Z., F.P., and H.K., generated and provided the ST data. M.S., L.W., S.K., H.K., and Y.Y., provided expertise and advised on data interpretation. W.W. conceived the study, planned and supervised the work, performed the analysis, and wrote the manuscript collaborating with all the other authors. All authors contributed to the interpretation of results and commented on and approved the final manuscript.

## Supplementary Figures

**Supplementary Figure S1.**
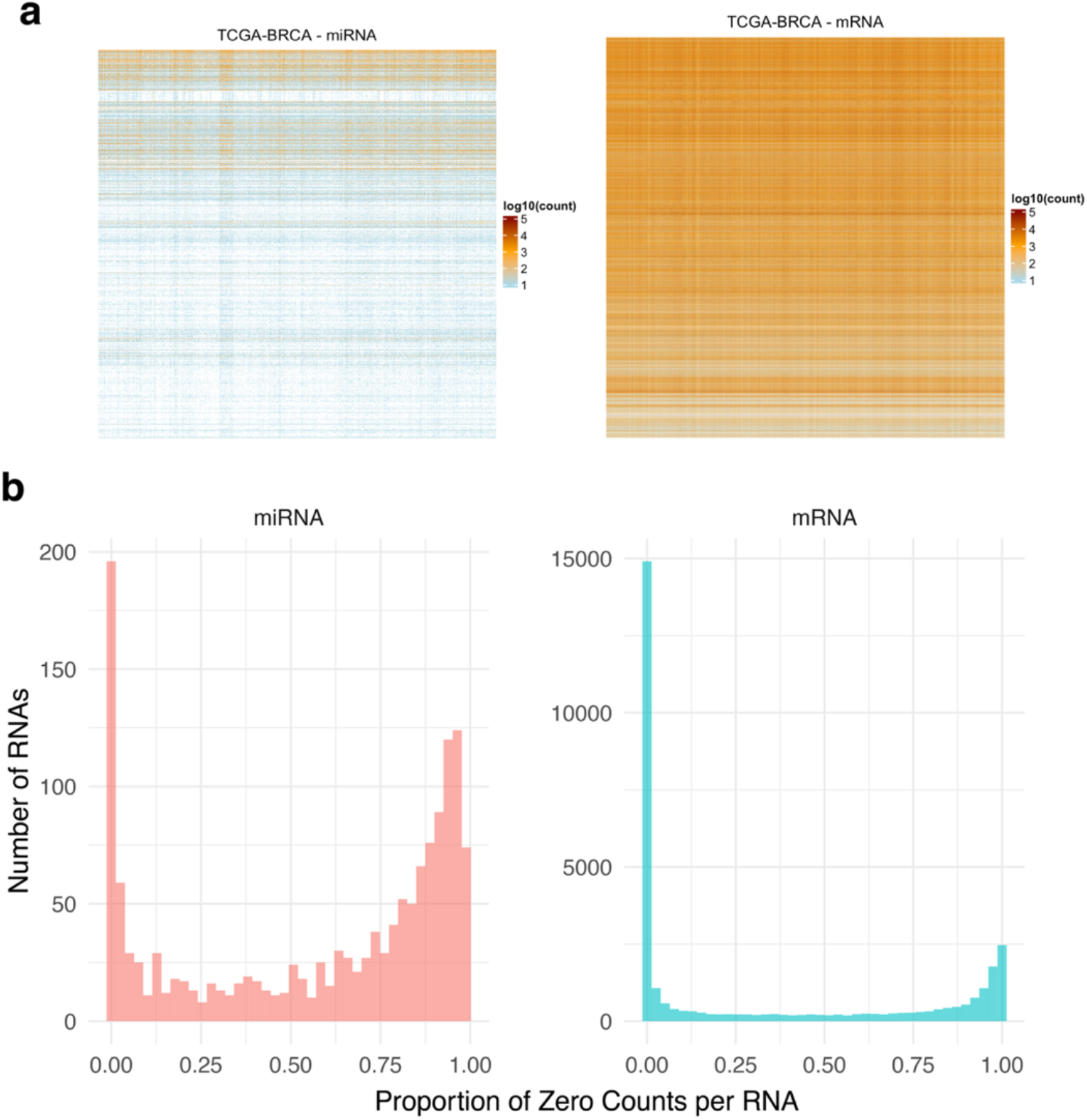
Comparison of expression count data between miRNA and mRNA in TCGA-BRCA samples. Unlike mRNA expression, which typically shows more continuous and log-normally distributed expression values, miRNA expression is characterized by extreme sparsity, with many miRNAs showing zero or near-zero expression across samples. (a) log_10_-transformed expression count matrices for miRNAs (left) and mRNAs (right). Each row corresponds to a gene or miRNA, and each column to a tumor sample. Zero counts are represented as white in the heatmap. (b) Distribution of the proportion of zero counts per gene across all samples. While most mRNAs are detected in nearly all samples, a large proportion of miRNAs are expressed in only a subset of tumors, with many exhibiting near-complete dropout.

**Supplementary Figure S2.**
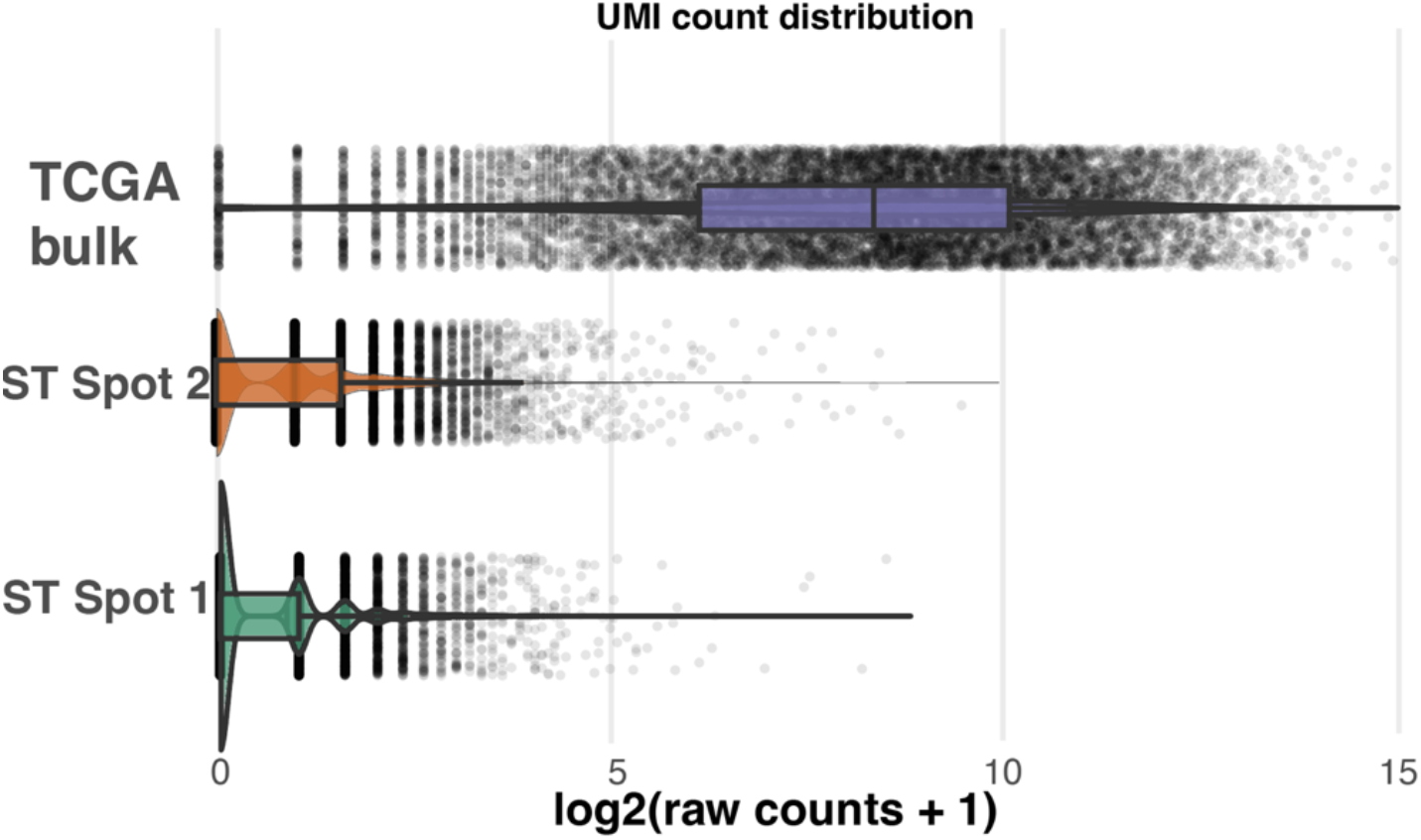
Comparison of expression sparsity between bulk RNA-seq in TCGA and random spatial transcriptomic (ST) spots. Distribution of *log*_2_ (raw counts+1) for genes measured in a random sample from TCGA breast cancer bulk RNA-seq and two randomly selected spots from a lung cancer ST sample. Each dot represents the raw count or UMI of one gene. Boxplots show interquartile range of gene expression levels with median indicated by the horizontal lines. The TCGA bulk RNA-seq profile exhibits a dense and continuous distribution, with a wide interquartile range, indicating effective log transformation and relatively low sparsity. In contrast, both ST Spot 1 and Spot 2 display highly compressed distributions near zero with extended right tails, indicating a high sparsity characteristic of ST data where most genes exhibit near-zero expression levels and can’t be modeled by log-normal distributions.

**Supplementary Figure S3.**
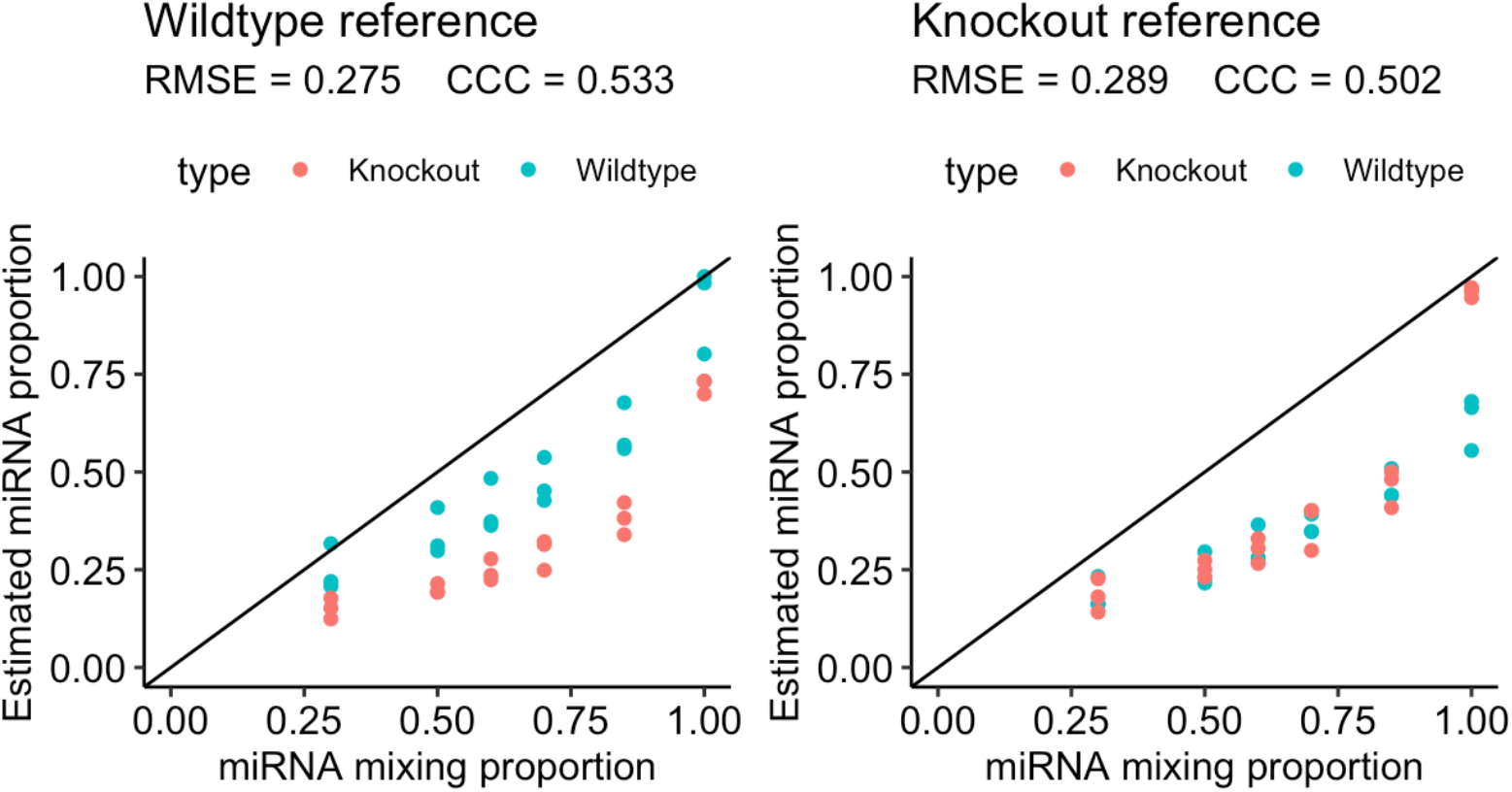
Comparison of DeconmiR performance when using *Dicer1* wildtype (left) or *Dicer1* knockout (right) as reference for all mixed samples.

**Supplementary Figure S4.**
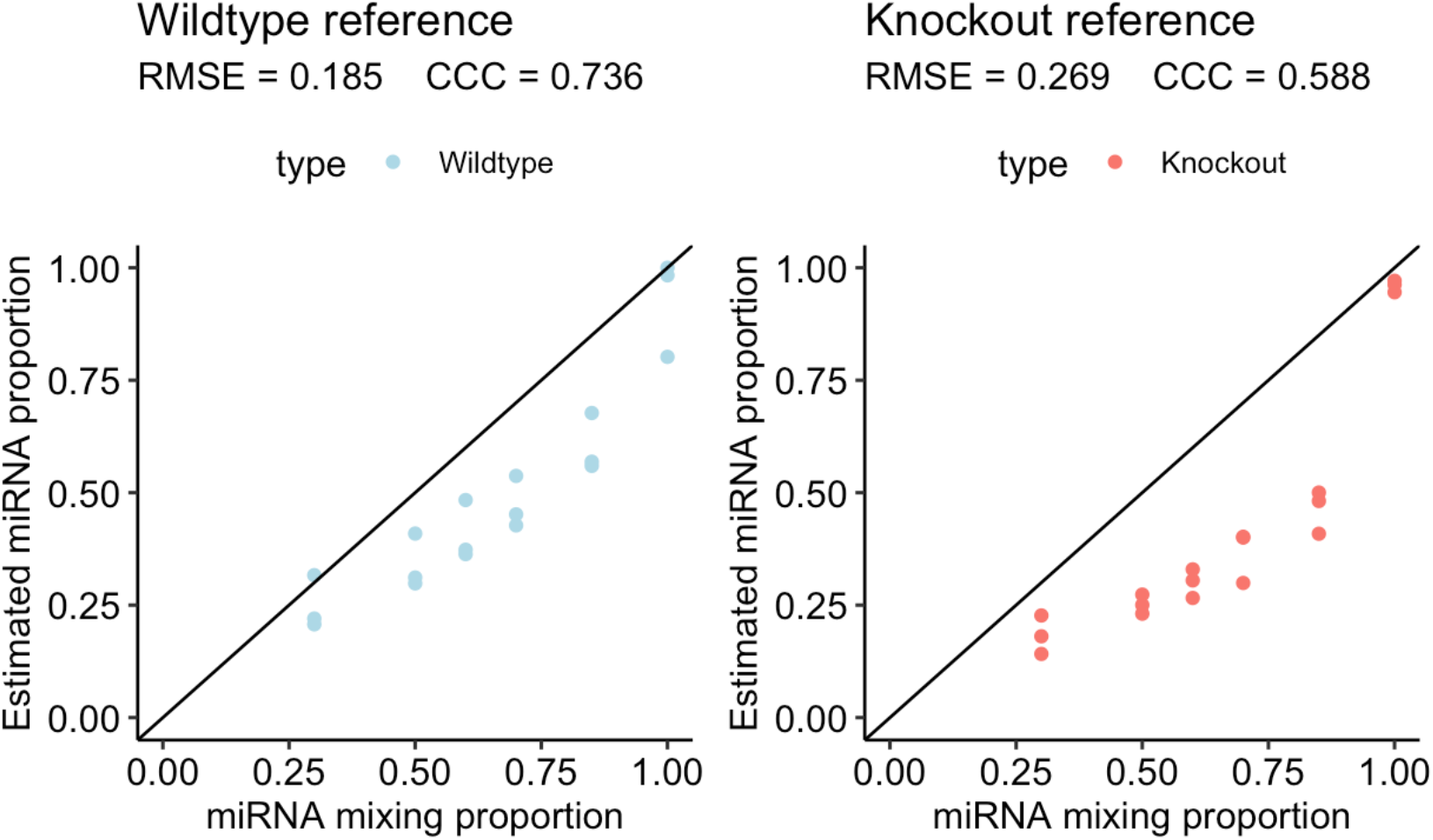
Comparison of DeconmiR performance when using *Dicer1* wildtype to deconvolve wildtype mixtures (left) or *Dicer1* knockout to deconvolve knockout mixtures (right). This deconvolution procedure requires knowing a priori which mixed samples are based on *Dicer1* wildtype and which ones are based on *Dicer1* knockout.

**Supplementary Figure S5.**
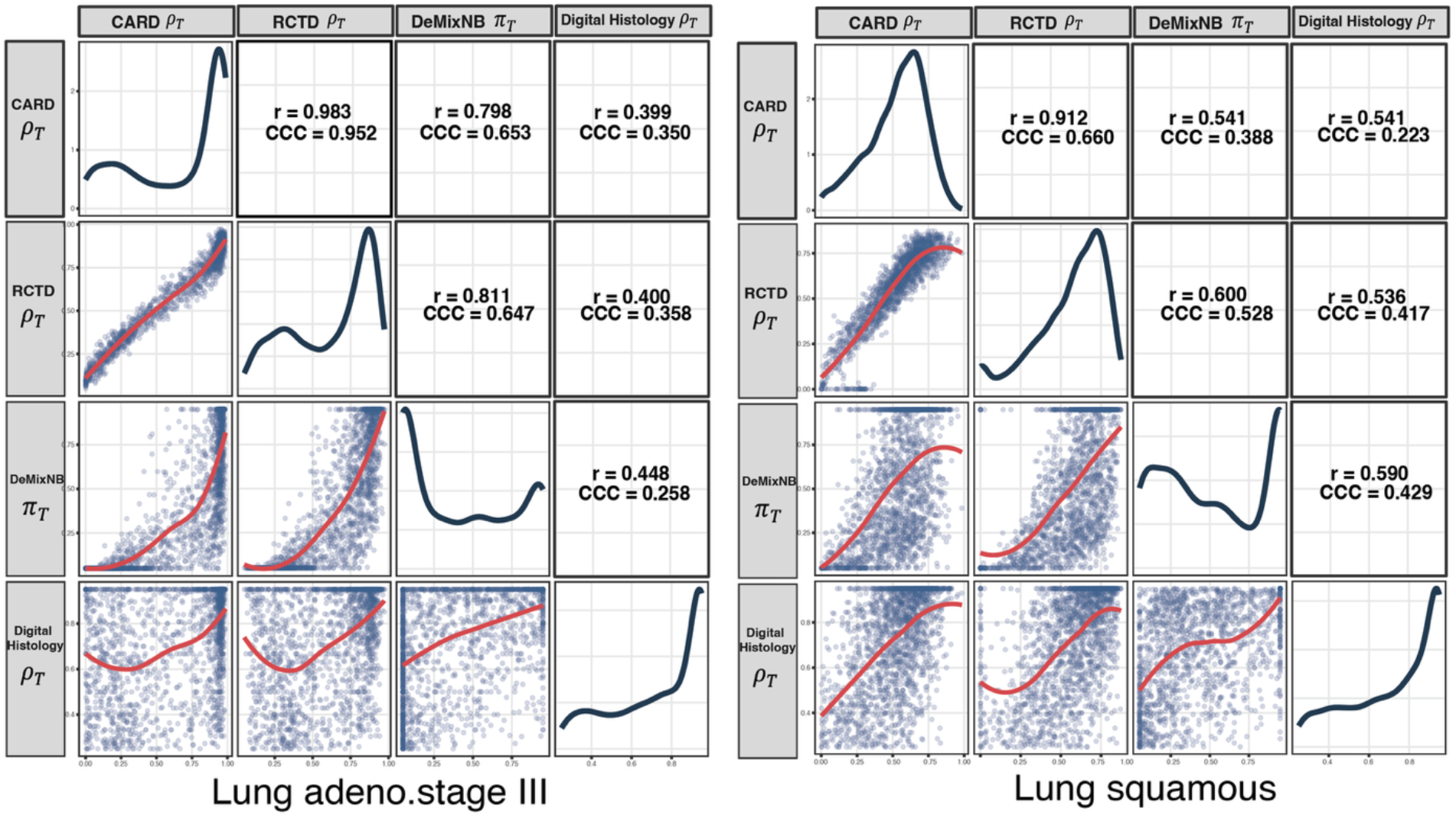
Pairwise comparison of tumor cell count proportions and tumor cell transcript proportions from deconvolution and digital pathology methods for a LUAD sample (left) and a LUSC sample (right). Pairwise scatter plot matrix comparing estimates from *ρ*_*T,CARD*_, *ρ*_*T,RCTD*_, *ρ*_*T,Digital Histology*_ and *π*_*T*_. The lower-diagonal panels show pairwise scatter plots with lowess() smoothing curves (red). The upper-diagonal panels display the Pearson correlation coefficients (r) and the Concordance Correlation Coefficient (CCC). The diagonal panels show the probability density distribution of estimates for each respective method. The moderate correlation demonstrated by *ρ*_*T,Digital Histology*_ and *π*_*T*_ between 0.45 and 0.59 matches what is observed in our miRNA-seq deconvolution study. On the other hand, even though *ρ*_*T,CARD*_ and *ρ*_*T,RCTD*_ are strongly correlated with each other, their correlations with *π*_*T*_ are at a moderate level for LUSC with balanced deviations on both sides of the smoothed curve. This is not the case for LUAD. Importantly in LUAD, *ρ*_*T,CARD*_ and *ρ*_*T,RCTD*_ identified a lot more spots as almost 100% tumor cells whereas *ρ*_*T,Digital Histology*_ reported a wide range of proportions. Hence we investigated these spots in Supplementary Figure S6.

**Supplementary Figure S6.**
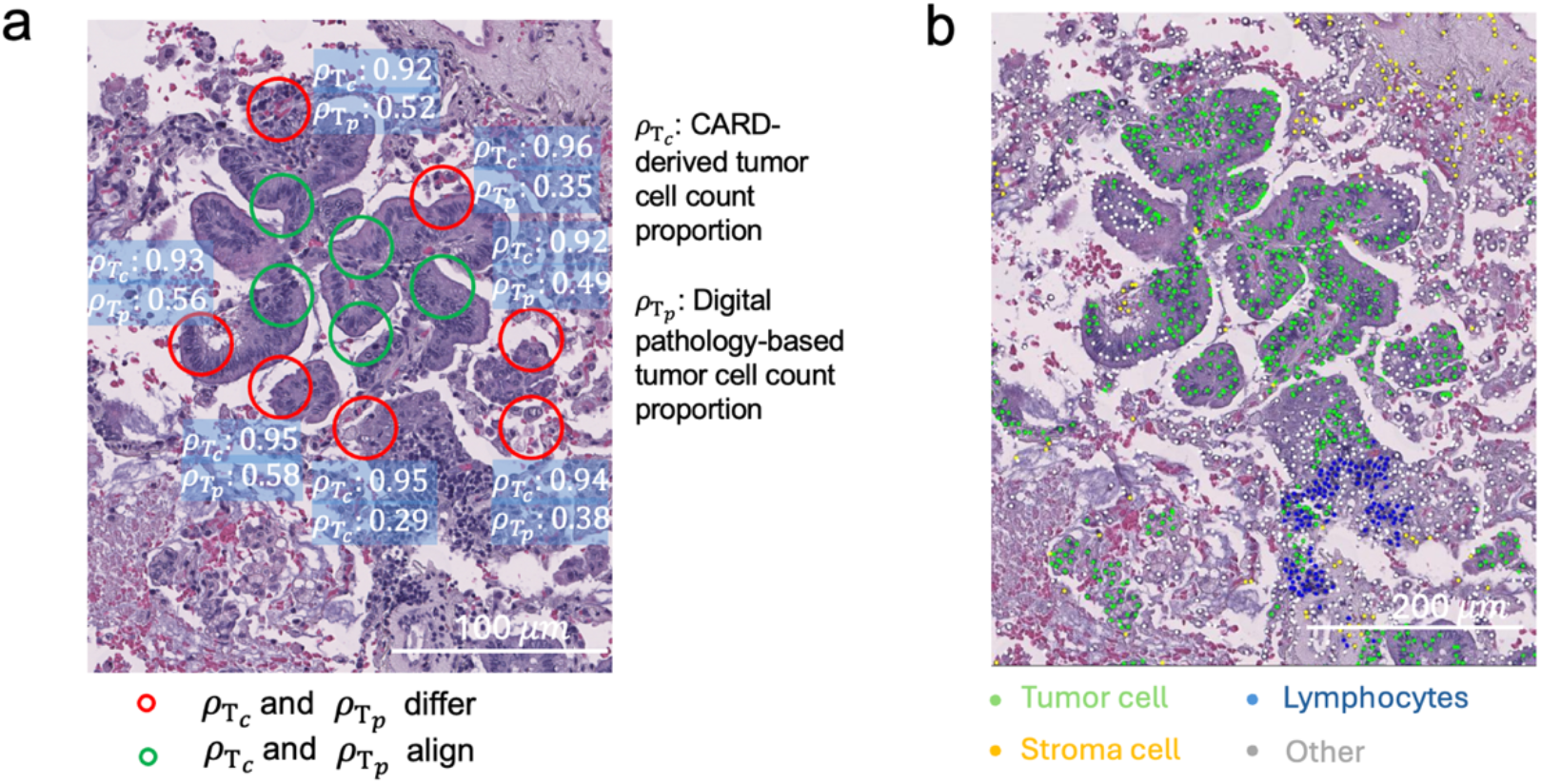
Validation of digital pathology-based tumor cell count proportions versus CARD in the lung adenocarcinoma stage III sample. (a) Representative region showing discordance between CARD-derived and digital pathology-based tumor cell count proportions. Green circles in the tumor core represent spots where the two methods align (difference < 0.1), red circles at the boundary represent spots where the two methods differ substantially (difference > 0.3). (b) Cell-type annotation of the same region showing tumor cells (green), lymphocytes (blue), stromal cells (yellow), and other cell types (gray). Digital pathology estimates 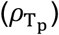 showed better consistency with the observed histological patterns, particularly in boundary regions with mixed cell populations.

**Supplementary Figure S7.**
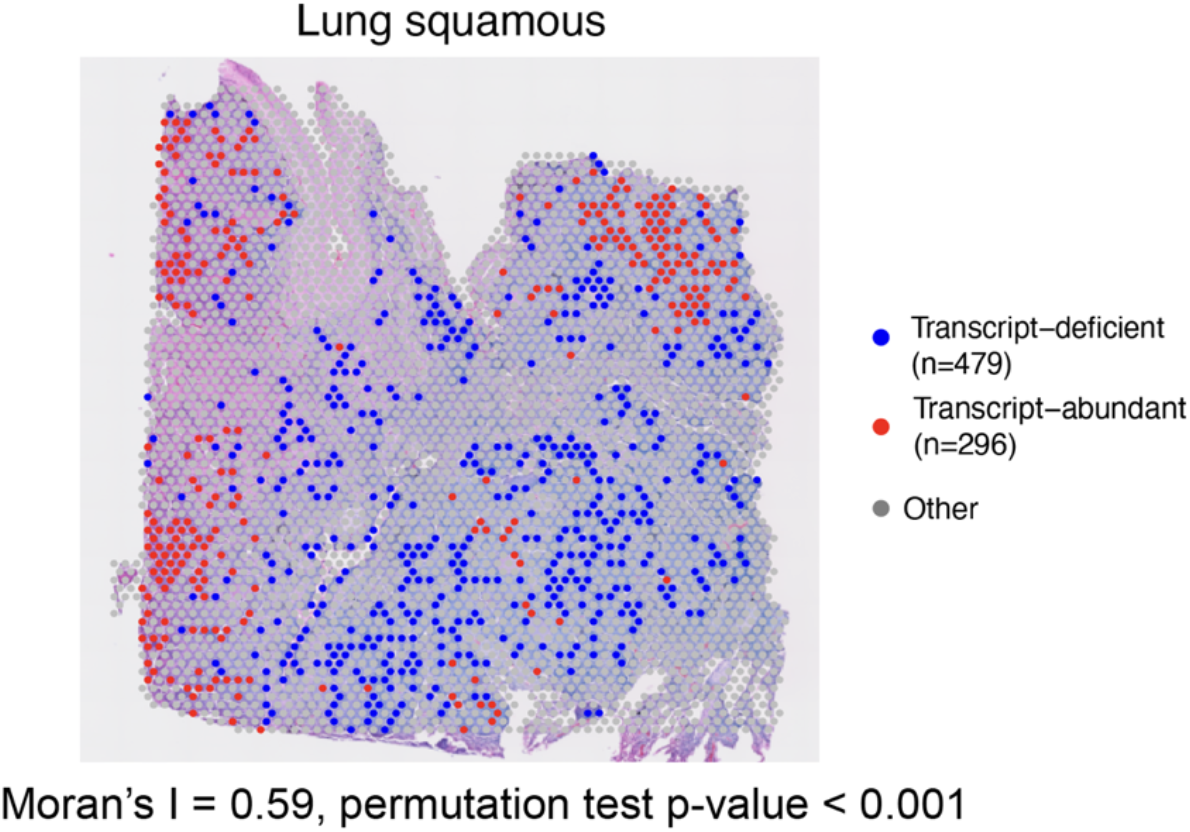
Spatial clustering of transcript-deficient and transcript-abundant spots for the lung squamous sample. Spatial visualization of transcript-deficient (blue) and transcript-abundant (red) spots overlaid on H&E histology image for lung squamous sample. Moran’s I spatial autocorrelation analysis demonstrates significant clustering (I=0.59, permutation test p<0.001).

**Supplementary Figure S8.**
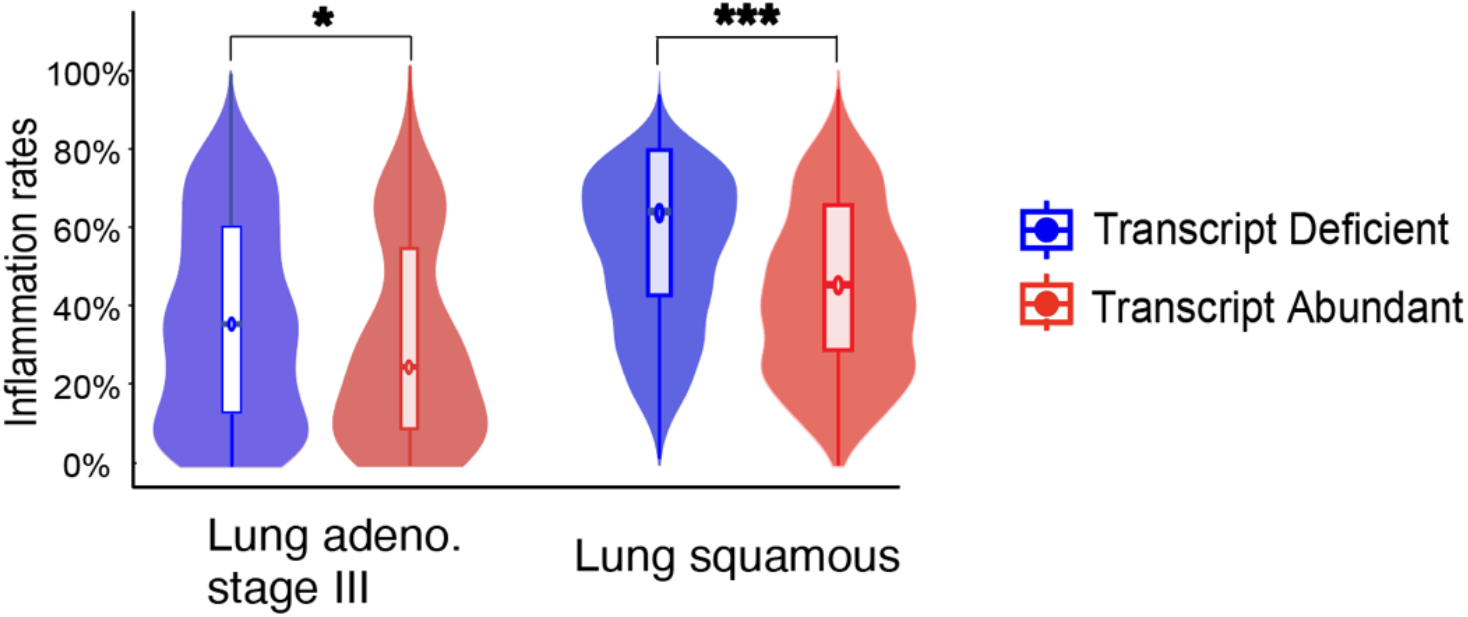
Comparison of local inflammation rates between transcript-deficient and transcript-abundant spots selected using an alternative difference threshold of 0.3. Inflammation rates represent the proportion of lymphocytes relative to the total lymphocyte and stromal cell count for each spot and its six adjacent neighbors. Lowering the threshold from 0.4 to 0.3 increased the number of transcript-deficient spots, replicating our fidings in both samples. Statistical significance was assessed using Wilcoxon rank-sum test with Benjamini-Hochberg (BH) correction. Significance level is denoted by: *, p < 0.05; **, p < 0.01; ***, p < 0.001; ****, p < 0.0001.

**Supplementary Figure S9.**
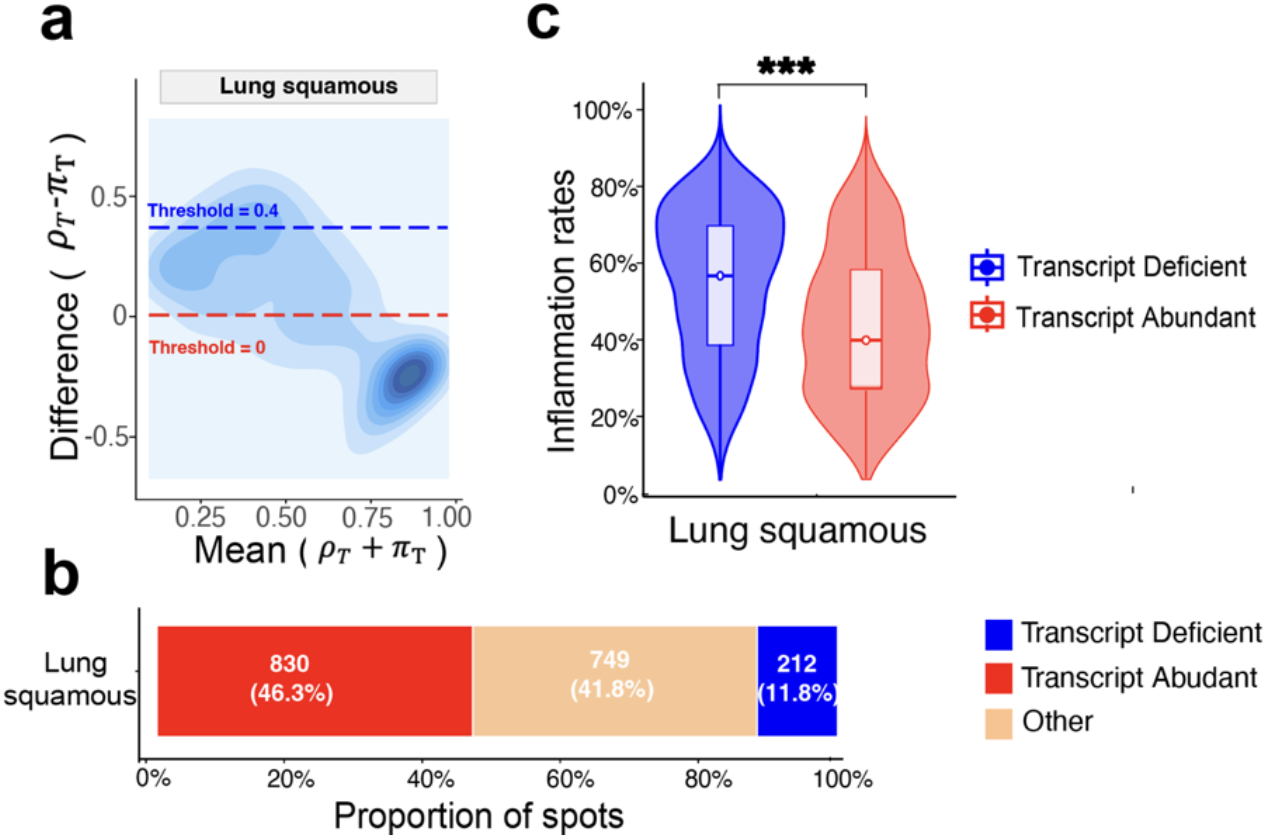
Validation of DeMixNB findings using CARD-derived tumor cell count proportions for the lung squamous sample. **(a)** MA density plot showing the distribution of differences (*D* = *π*_*T*_ − *ρ*_*T*_) between tumor transcript proportions and CARD-derived tumor cell count proportions versus the means. Horizontal dashed lines indicate thresholds for categorizing transcript-deficient (*D* ≥ 0.4, blue) and transcript-abundant (*D* ≤ 0; red) spots. **(b)** Bar plot shows the proportion of spots in each category. **(c)** Comparison of local inflammation rates between transcript-deficient and transcript-abundant spots. Inflammation rates represent the proportion of lymphocytes relative to the total lymphocyte and stromal cell count for each spot and its six adjacent neighbors. Statistical significance was assessed using Wilcoxon rank-sum test with Benjamini-Hochberg (BH) correction. Significance level is denoted by: *, p < 0.05; **, p < 0.01; ***, p < 0.001; ****, p < 0.0001.

